# CyanoTag: Discovery of protein function facilitated by high-throughput endogenous tagging in a photosynthetic prokaryote

**DOI:** 10.1101/2024.02.28.582475

**Authors:** Abigail J. Perrin, Matthew Dowson, Adam A. Dowle, Grant Calder, Victoria J. Springthorpe, Guoyan Zhao, Luke C. M. Mackinder

**Author notes:** Corresponding authors &.

## Abstract

Despite their fundamental importance to aquatic ecosystems, global carbon cycling and exciting potential in sustainable bioindustries, the genomes of photosynthetic prokaryotes still contain large numbers of uncharacterised protein-coding genes. Here we present a high-throughput approach for scarless endogenous fluorescent protein tagging in the model cyanobacterium *Synechococcus elongatus* PCC7942. From 400 targets we successfully endogenously tag over 330 proteins corresponding to >10% of the proteome. We demonstrate how this resource can be used at scale to determine subcellular localisation, track relative protein abundances and to elucidate protein-protein interaction networks. Our data has provided biological insights into a diverse range of processes - from photosynthesis to cell division. Of particular interest, our ‘CyanoTag’ lines enabled us to visualise in real time the rapid condensation of Calvin Cycle proteins Prk and Gap2 within seconds of withdrawal of light, effectively ‘switching off’ photosynthesis in the dark. These insights, CyanoTag cell lines, associated data and optimised methods are intended to be shared as a resource to facilitate further discoveries relevant to cyanobacteria and more broadly to all photosynthetic life.

## Introduction

Cyanobacteria fulfil essential roles in maintaining Earth’s atmosphere and ecosystems. These photosynthetic prokaryotes make a significant contribution to global CO_2_ fixation, O_2_ production and nutrient cycling^1,2^. They form intricate connections and symbioses across diverse ecosystems with their imbalance leading to disastrous effects^3^. Via ancient primary endosymbiosis events that led to the development of chloroplasts in eukaryotes, cyanobacteria have shaped the evolution of hugely diverse plant and algal species^4^. Many photosynthetic genes and central metabolic processes remain conserved between cyanobacteria and eukaryotic chloroplasts. Thus, understanding cyanobacterial cell biology can contribute to advances in knowledge of fundamental cellular processes, chloroplast biology and endeavours in photosynthetic engineering of plants^5,6^. In addition, their diversity, genetic tractability and photosynthetic capability position cyanobacteria as promising chassis for green biotechnology applications, including carbon capture and high-value product manufacture^7^.

*Synechococcus elongatus* PCC7942 (*S. elongatus* hereafter) is a freshwater cyanobacterium, the first to be transformed with exogenous DNA^8^ and now a commonly-used model organism. Its compact (∼2.7 Mbp) genome and elongated morphology with identifiable cellular structures also make it an attractive organism for study. Despite the global importance of cyanobacteria and their widespread use in research, a strikingly small proportion of cyanobacterial proteins have been characterised; over a quarter of *S. elongatus* genes lack any functional annotation, with most of the remainder annotated only via bioinformatic methods (through genetic homology to characterised genes in other species). Hence there are significant gaps remaining in our understanding of fundamental biology relating to photosynthesis, cell division, central metabolism and more.

The use of large-scale tagging approaches has led to significant insights in model systems^9^. Most studies to date have focused on model organisms including yeast^10,11^, *Caenorhabditis elegans*^12^, drosophila^13^, and human cells^14^. Some more recent studies have started to develop and apply large-scale localisation approaches to other eukaryotic groups including the parasite *Trypanosoma brucei*^15^ and the photosynthetic alga *Chlamydomonas reinhardtii*^16,17^. Outside of *Escherichia coli*^18^ large-scale tagging has had limited application in prokaryotic systems and has yet to be applied to a photosynthetic prokaryote. Furthermore, the application of endogenous, scarless fluorescent protein tagging via chromosomal integration and subsequent removal of markers has yet to be applied at scale to photosynthetic systems. To enable the rapid advancement of our knowledge of photosynthesis and cyanobacterial cell biology, we built a pipeline for the generation of a library of *S. elongatus* lines expressing fluorescently-tagged proteins amenable for use in protein localisation, expression and interaction studies. Our approach involves the scarless tagging of each gene at its native locus, maintaining endogenous regulatory regions and allowing tag fluorescence to be used as a proxy for protein abundance.

Here we report the tagging and further characterisation of 330 protein-coding genes (around 12% of the *S. elongatus* genome), and examples of novel biological insights this has already provided. We also present a preliminary protein interactome based on affinity purification experiments where we detected over half the 2,714 known proteins in this organism. The interactome map is the first of its kind for a cyanobacterial species and includes 369 high-confidence protein-protein interactions. As we expand our libraries towards coverage of the entire *S. elongatus* proteome, we are sharing the growing datasets via an interactive web tool (https://morf-db.org/projects/York-Mackinder-Lab/MORF000032). We anticipate our cell libraries, associated data and optimised methods (see Supplementary information) being valuable resources for research communities, providing insights into both known processes and uncharacterised proteins in cyanobacteria and beyond.

## Results

### Development of CyanoTag, a high throughput protein tagging platform

To advance our understanding of the *S. elongatus* proteome, we developed an approach to scarlessly tag proteins at their native C-termini in a way that would facilitate the determination of each protein’s subcellular localisation, relative abundance and interacting proteins (Fig. 1a). We used a modified Golden Gate cloning approach to construct plasmids (Fig. 1b), such that the constructs’ integration into the genome by homologous recombination results in the 3’ tagging of the gene with the bright monomeric green fluorescent protein mNeonGreen (mNG) and a 3xFLAG epitope, as well as the introduction of positive and negative selectable markers downstream of the gene of interest and flanked by copies of mNG to enable subsequent marker removal via a second homologous recombination event (Fig. 1b).

**Fig. 1:**
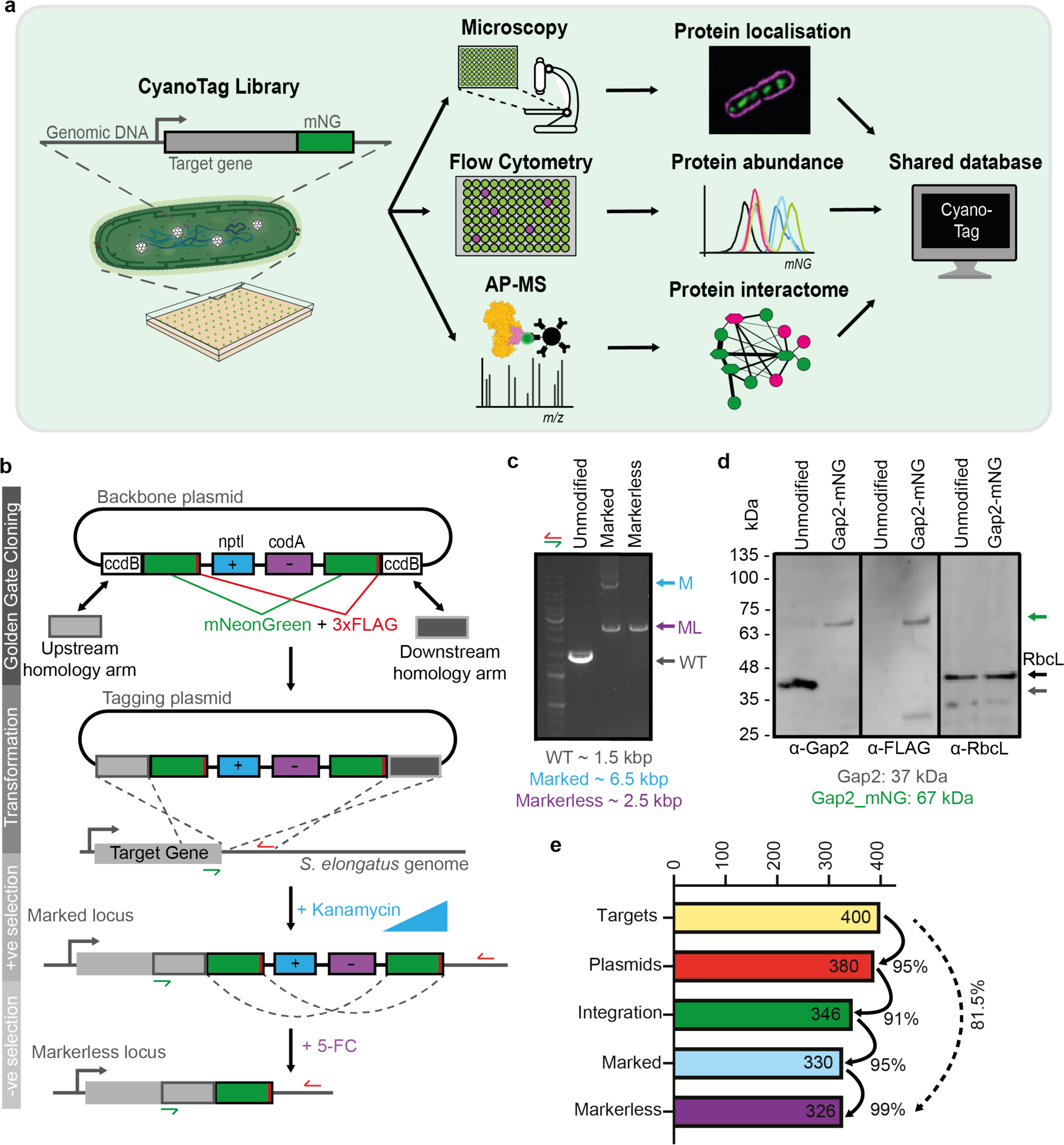
A high-throughput fluorescent protein tagging pipeline in *S. elongatus* PCC 7942. **a,** Overview of the multipurpose 96-well pipeline, which can be used to tag proteins of interest at their native loci for the study of protein distribution, protein abundance and protein-protein interactions. AP-MS: affinity purification-mass spectrometry. **b,** Schematic showing the pipeline for high-throughput protein tagging. Two homology arms (HAs) are amplified by PCR from genomic DNA, and a BsaI- or BspQI-dependent Golden Gate cloning reaction is used to insert these HAs in place of two cytotoxin-encoding (*ccdB*) genes in a backbone plasmid which contains a series of tagging and selection modules. These are an mNeonGreen fluorescent protein and 3xFLAG epitope (mNG/3xFLAG) tagging module, a kanamycin antibiotic selection marker (*nptI*) and a negative selection marker (*codA*) followed by a second copy of the mNG/3xFLAG module. Following transformation of *S. elongatus* PCC7942 with these plasmids and selection with kanamycin, homologous recombination leads to integration of the tagging and selection modules at the 3’ end of the target gene and the generation of marked mutants. Markerless mutants are generated via a second recombination event, enhanced by selection with 5-fluorocytosine (5-FC), between the two mNG/3xFLAG modules. **c,** Example of colony PCRs showing the changes in locus size as homoplasmic marked and markerless mutants are generated in the CyanoTag pipeline. We observe a degree of ‘spontaneous’ marker removal before the addition of 5-FC in most marked lines. Primer locations are indicated with half-arrows on (b). **d,** Western blots showing the modification of the Gap2 protein (*Synpcc7942_1742*) upon tagging. Blots for Rubisco large subunit (*Synpcc7942_1426*) are shown as a loading control. **e,** Bar graph showing the efficiency at each step shown in (b) and the overall success rate of the pipeline.

We transformed *S. elongatus* with each plasmid individually and selected for transgene integration based on kanamycin resistance, conferred through the positive selection marker *nptI.* As *S. elongatus* is polyploid we used increasing concentrations of kanamycin to ultimately isolate homoplasmic “marked” mutants, in which every chromosomal copy of the gene of interest was modified. To remove both selection markers to create “markerless” tagged mutants, we induced the excision of the selectable markers via a second round of homologous recombination of the flanking mNG genes using 5-fluorocytosine (5-FC) which is hydrolysed to cytotoxic 5-fluorouracil by the *codA* gene product (negative selection). We tested our pipeline on 400 target genes primarily associated with photosynthesis and other processes of particular interest to the cyanobacteria research community, using 96-well plate formats for cloning, transformation and selection steps. Homoplasmic integration of the tagging construct and successful marker removal were validated in each case by colony PCR (Fig. 1c) and introduction of the mNG and FLAG tags caused an ∼30 kDa increase in observed protein molecular weight (Fig, 1d).

Each step in the plasmid cloning and selection of marked mutants had a success rate of between 91% and 99%, allowing us to isolate homoplasmic tagged mutants for over 330 (82.5%) of our target genes. (Fig. 1e). Our tagging success rate is comparable to other large-scale endogenous tagging studies^11,14,15^ and higher than the ∼53% exogenous tagging success rate in the photosynthetic alga *C. reinhardtii* that faces significant gene cloning challenges due to intron essentiality and high GC content^16,17^.

### Protein localisation in markerless mutants

We used super-resolution structured illumination microscopy (SIM) to observe mNeonGreen fluorescence in 322 markerless mutants. To aid classification of the fluorescent protein distribution for each tagged cyanobacterial strain we built a decision tree based on the pattern of mNG fluorescence and how it relates to the consistent pattern of cellular autofluorescence, which primarily arises from the pigments in the thylakoid membranes (Fig. 2a). We used this to describe the patterns of fluorescence observed in each line using one or a combination of descriptors relating to diffuse patterns (D), membrane patterns (M) or puncta (P) (Fig. 2b, 3a, 3b & S1).

**Fig. 2:**
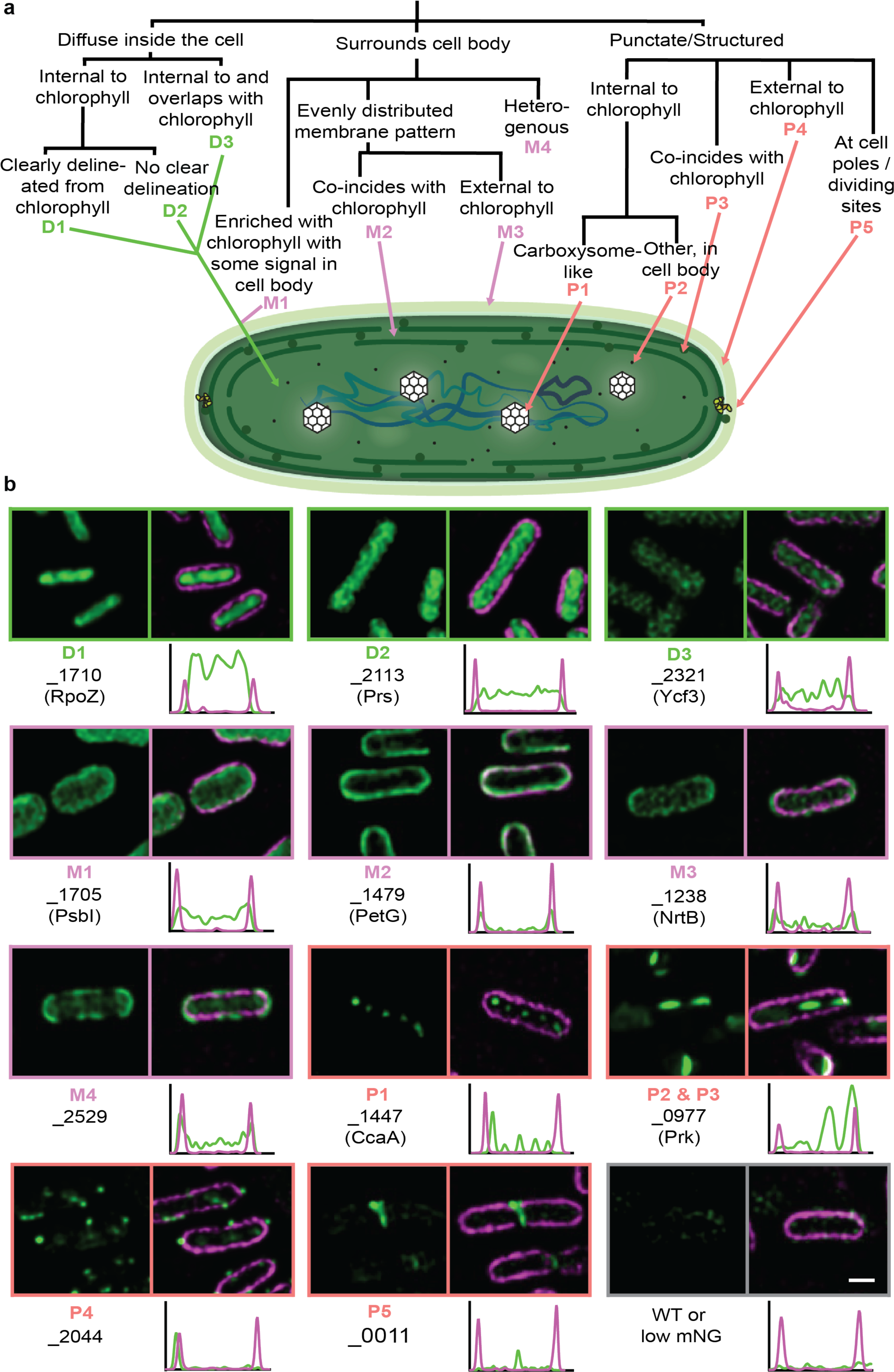
Subcellular locations of tagged proteins in *S. elongatus PCC 7942*. **a,** Schematic overview of a decision tree for assignment of proteins to classes (or combinations of classes) according to the localisation of mNG fluorescence. **b,** Representative examples of tagged proteins (green) localising to each cluster in the set of *S. elongatus* markerless mutants. Chlorophyll autofluorescence signals (magenta) are overlain in the right hand panels. Underneath each overlay the profile of fluorescence intensity across the length of the displayed cell is represented. The transects and raw values from these analyses are displayed in Fig S1. Scale bar: 1 μm.

**Fig 3.**
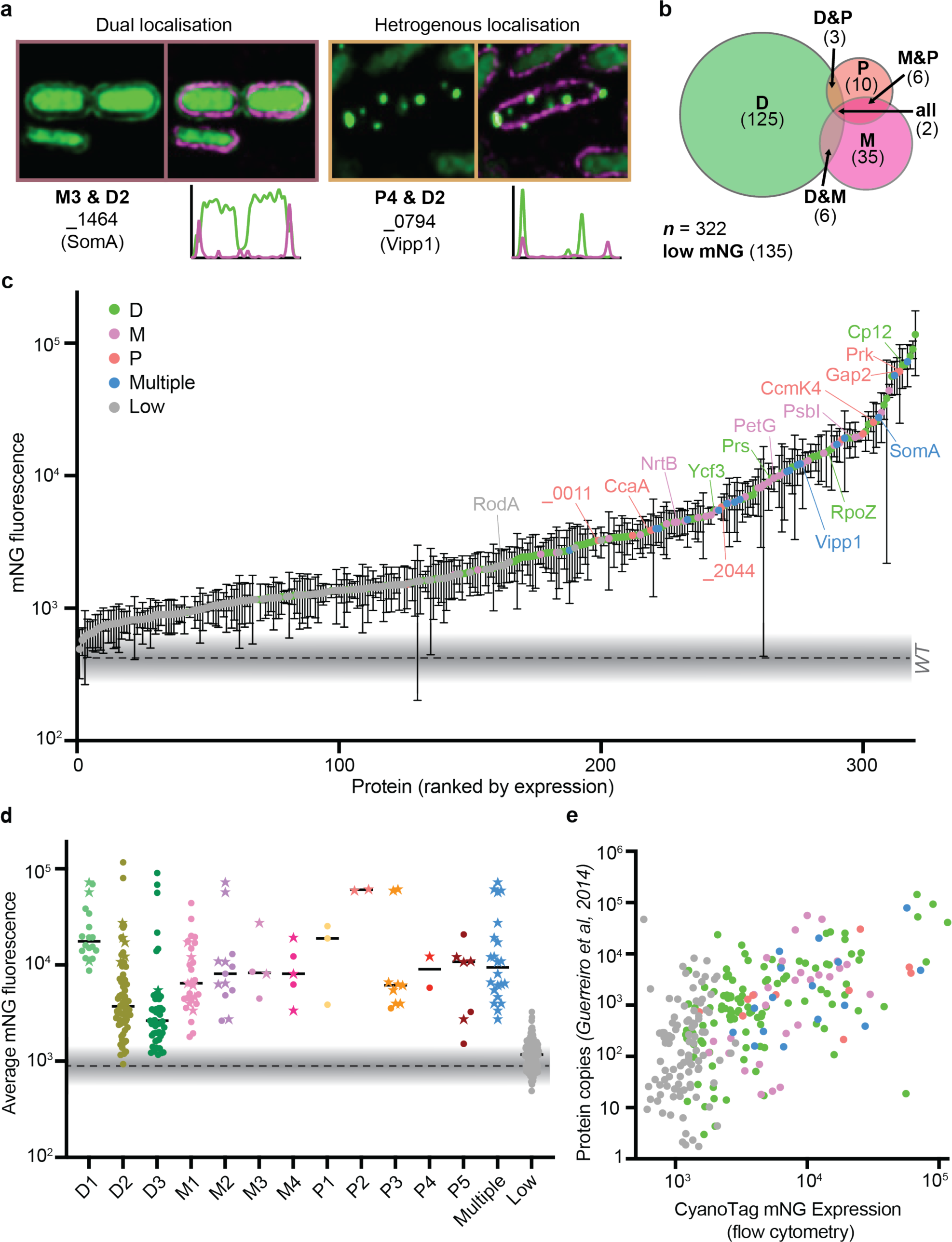
CyanoTag lines facilitate interrogation of protein expression as well as localisation in live cells. **a,** Example images of lines with multiple expression locations (see Fig 2), either within the same cell (left) or with different patterns in different cells. **b,** Venn diagram summarising broad localisation descriptors, and the crossover between them, for the imaged lines. **c,** Normalised fluorescence intensities for markerless lines. Means and standard deviations for 3 or more replicates are plotted. **d,** Mean fluorescence values of markerless lines, by localisation category. Star shaped points indicate that multiple localisation descriptors are used. **e,** Scatter plot of estimated protein copy number per cell, (averaged across a 48 hr time course of 12:12 hr light/dark cycles from a previously published study) plotted against mean fluorescence intensity recorded in corresponding markerless CyanoTag line.

Amongst these images we observe a range of distinctive and/or novel localisation patterns. For instance, using three lines where known carboxysome proteins were tagged, we see a characteristic punctate pattern of mNG fluorescence along the central axis of the cell (P1) (Fig. 2b). In multiple other examples, we show previously undescribed localisations for poorly characterised cyanobacterial proteins such as Synpcc7942_2044 (P4) (Fig. 2b). Little is known about Synpcc7942_2044, beyond its likely essentiality^19^ and as containing a PRC-barrel (Photosynthetic Reaction Center-barrel), which are speculated to have been acquired from purple bacteria and incorporated into the regulation of cyanobacterial electron transport^20^. Its localisation to distinct puncta external to the thylakoids hints either to a novel regulatory mechanism or a different function entirely.

A small proportion (<9%) of the proteins we describe here do not fit neatly into a single localisation category (Fig 3a, 3b, S1). The Vipp1 protein (Synpcc7942_0794) is one such example, where under our imaging conditions (low-light) we see a disperse cytoplasmic localisation of tagged Vipp1 protein in most cells, but very bright puncta at the cell periphery in others (Fig. 3a). Based on previous observations, it is likely that in these cells Vipp1 is playing a role in maintaining the integrity of the thylakoid membrane and/or protecting the cell from stress-induced damage^21,22^.

### The CyanoTag pipeline enables screening of *in vivo* protein abundances

Due to the maintenance of both native *cis* and *trans* regulatory elements in the markerless lines, mNG can act as a strong proxy for protein abundance. Based on this property we developed a simple flow cytometry-based fluorescence assay (Fig. S2a) and used it to determine the relative abundance of proteins in live cells under standard growth conditions (Fig. 3c & 3d). Fluorescence intensity was a strong predictor that protein subcellular location could be assigned from imaging data (Fig. 3c & 3d). The trends in expression levels we observe in our flow cytometry data broadly reflect those observed in published mass spectrometry datasets (Fig. 3e)^23^. This assay provides a platform by which changes in protein expression, and also cellular morphology, can be assessed in a high-throughput manner (examples in later sections).

### The CyanoTag pipeline facilitates comprehensive mapping of protein interactions

To explore the protein-protein interaction network in this cyanobacterium, we developed an affinity purification-mass spectrometry (AP-MS) method and applied it in duplicate to an initial cohort of markerless CyanoTag lines (Fig. 4a, Table S2). We then developed a bioinformatic pipeline to filter mass spectrometry data, using a combination of CompPASS^24^ and SAINT^25^ analyses to define high confidence interactors (HCIs) for each of the 82 bait proteins (Fig. 4a & 4b). The data collated from these initial bait proteins resulted in the identification of two or more peptides corresponding to 1,617 unique prey proteins representing ∼60% the entire *S. elongatus* proteome (Fig. 4a). Of the 56,422 protein interactions detected, after filtering, 369 passed the thresholds to be HCIs. These data were compiled into an initial protein interactome comprising 82 baits and 135 unique prey proteins, which includes previously characterised interactions and highlights potentially novel ones (Fig. 4c & S3).

**Fig. 4:**
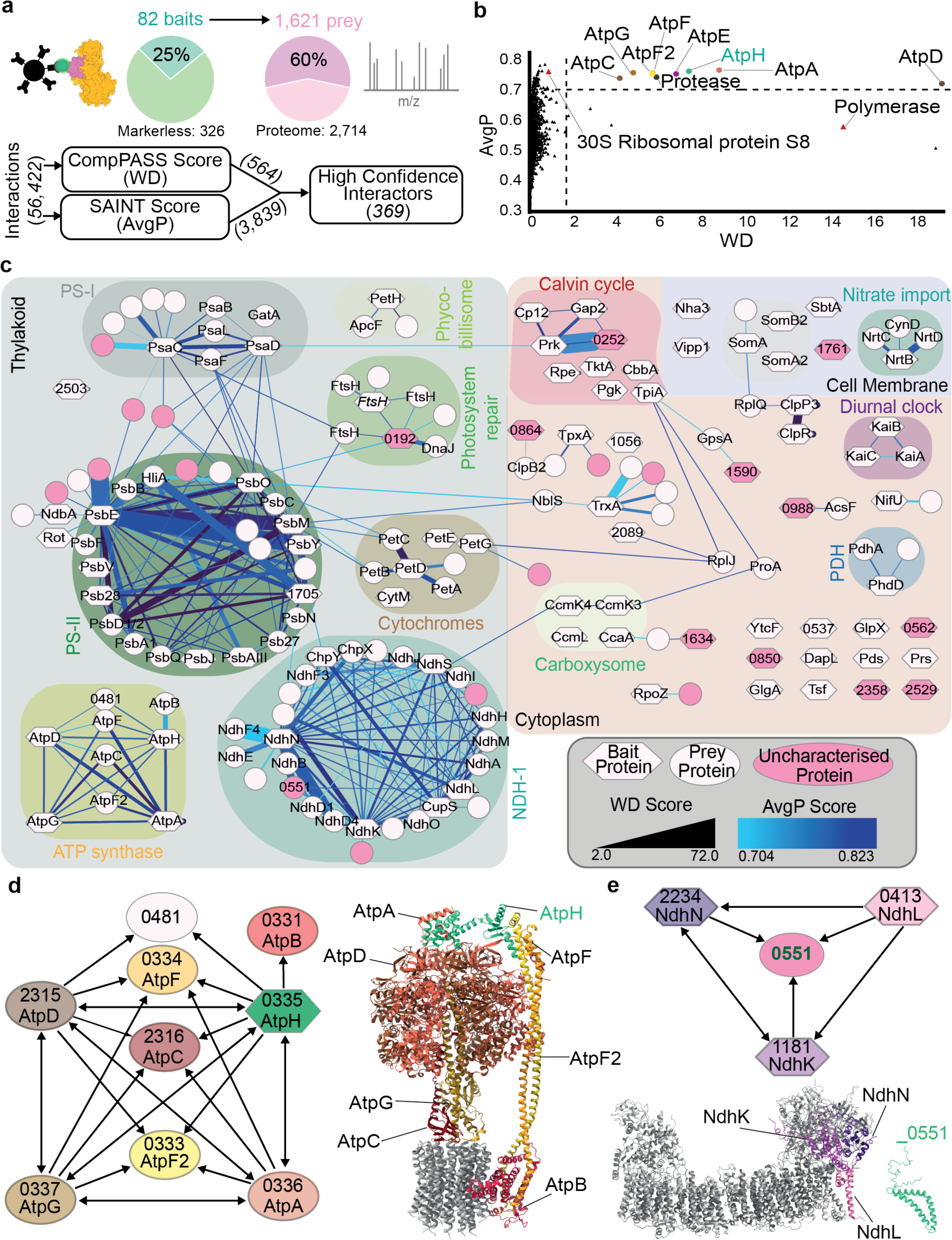
A preliminary *S. elongatus* protein interactome using CyanoTag lines. **a,** Overview of the AP-MS and bioinformatic filtering pipeline, which was used to identify high confidence protein interactions from a biologically diverse range of bait proteins. **b,** Representation of bioinformatic filtering of protein interactions for the bait AtpH, a well characterised component of the ATP synthase complex. Using just one confidence score (WD or AvgP) results in the presence of likely false positive interactions (red triangles). Combining both preserves almost all previously-characterised complex interactions (labelled). **c,** Preliminary protein interactome of *S. elongatus* based on data from 82 bait proteins run in duplicate. Labels on nodes denote proteins by an abbreviated name if one exists or by their four number gene identifier otherwise. Node shape corresponds to bait or prey status and edge width and colour represent WD and AvgP scores respectively. Known complexes are highlighted by background colour and currently uncharacterised protein nodes are shaded pink. **d,** Interactions of the bait protein AtpH (highlighted green) with its high confidence preys (left). Model of the ATP synthase complex (PDB: 6OQR) highlighting bait and prey proteins (right). **e,** Interactions of the prey protein Synpcc7942_0551 (green) and the three bait proteins that it co-purified with as high confidence interactors (NdhN, L and K) (top). AlphaFold2 predicted structure of Synpcc7942_0551 and model structure of the NDH1-MS complex (PDB: 6TJV) with interacting subunits highlighted (bottom).

The resulting interactome map includes multiple interactions within and between known large protein complexes, including photosystems I and II, the NDH-1 complex, nitrate import proteins and the circadian clock regulatory proteins KaiABC (Fig. 4c). In cyanobacteria the NDH-1 complex is found in four isoforms, all that share a core common module but have different isoform specific subunits^26^. Using the core structural component NdhN (Synpcc7942_2234) as bait eighteen other core or isoform specific NDH-1 subunits were identified as HCIs (Fig. 4c). Using the AtpH subunit (Synpcc7942_0335) of the ATP synthase complex as a bait, we detected HCIs with almost all of the known proteins of this highly conserved complex^27^ (Fig. 4b, 4c & 4d). Only subunits of the membrane integral C-ring were not classified as high-confidence interactors of AtpH. All four subunits of ATP synthase used as baits each identify interactions - almost exclusively - with multiple other subunits of the complex (Fig. 4c & 4d).

Amongst the HCIs are several examples of potentially novel interactions. Synpcc7942_0551 is an uncharacterised prey protein identified as interacting with three NDH1-MS complex subunits (Fig. 4c & 4e). The interacting NDH1 subunits are adjacent to each other and in close proximity to the plastoquinone-binding site of the complex (Fig. 4e). This suggests the AP-MS pipeline could be sufficiently powerful to resolve novel interactions to specific regions within a protein complex. Synpcc7942_0551 appears to be conserved across many cyanobacteria species and is structurally consistent with being a transmembrane protein, however is absent in *Thermosynechococcus elongatus* BP-1, the thermophilic cyanobacterium used for structural determination of the of the NDH1 complex^28,29^.

### Analysis of CyanoTag lines provides insights into the regulation of the Calvin cycle

In the course of developing the CyanoTag pipeline, we observed many striking or unexpected protein localisation patterns in our fluorescently tagged lines, as well as evidence for novel protein-protein interactions. Amongst these were observations related to the regulation of the Calvin cycle, where we identified Synpcc7942_0252 as a potential novel interactor of Gap2 and Prk (Fig. 4c & 5a). This protein contains a Cp12-like domain but appears from our imaging and expression data to be present at a level much lower than Cp12 (Fig. 5b & 5c). Using our flow cytometry method, alongside fluorescence microscopy, we detected increased fluorescence of mNG-tagged Synpcc7942_0252 after several hours of darkness (Fig. 5c & 5d), corresponding with published analysis that showed the transcription of the corresponding gene to be amongst the most highly upregulated in response to a two-hour dark pulse^30^.

**Fig. 5:**
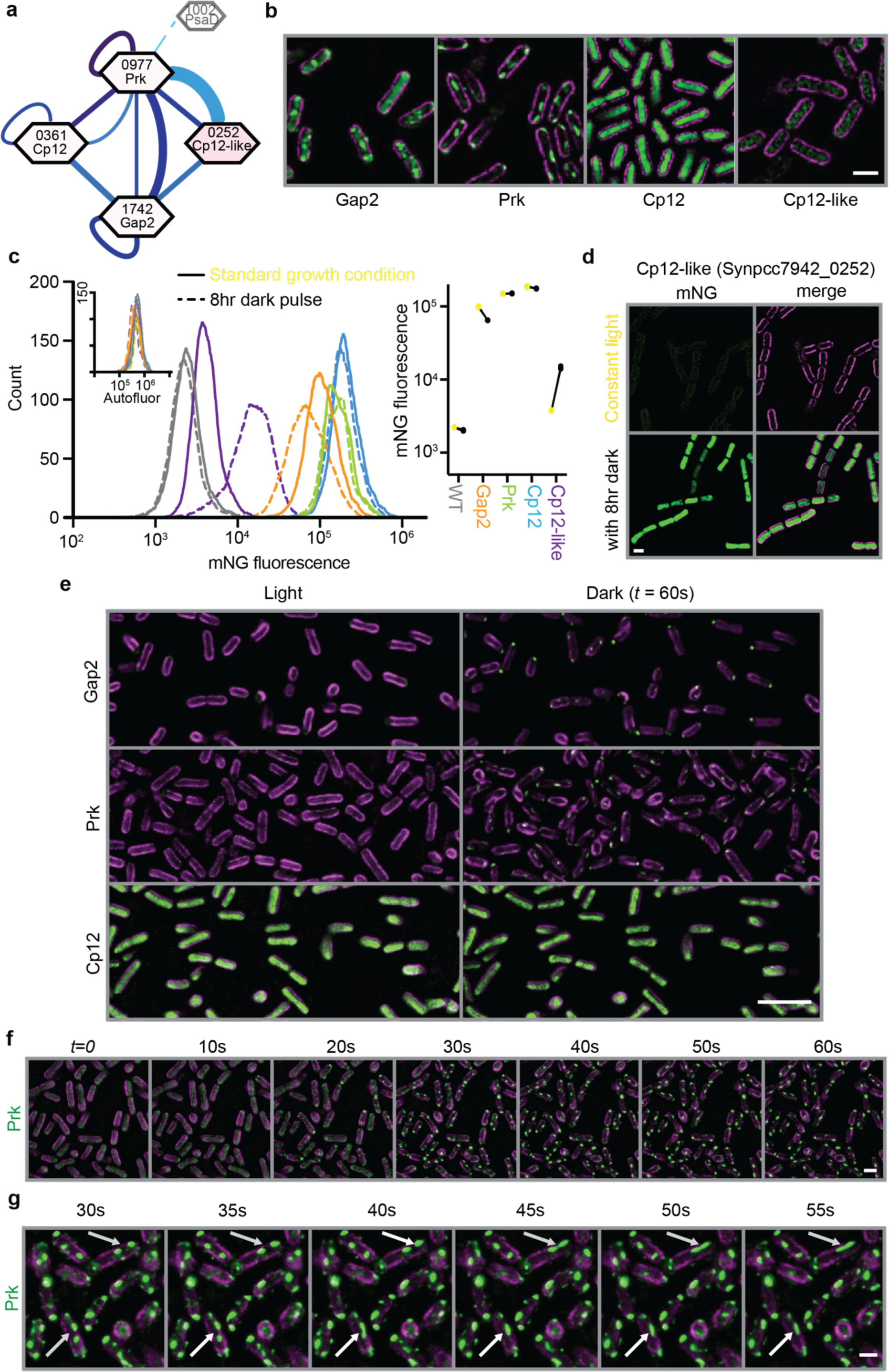
CyanoTag mutants allow visualisation and investigation of light-dependent regulation of the Calvin cycle. **a,** Interactions detected between Gap2, Prk, Cp12 and Cp12-like (Synpcc7942_0252) by our AP-MS pipeline. **b,** Images of Gap2, Prk, Cp12 and Cp12-like collected under standard conditions (grown in constant light, then exposed to at least 5 minutes of darkness in the imaging chamber). **c,** Flow cytometry-based quantitation of the mNG fluorescence of these proteins with and without 8 hours incubation in the dark. Left: Histograms of mNG fluorescence detected in a representative sample of each. Inset are the corresponding histograms showing the same samples’ autofluorescence (Allophycocyanin channel). Right: median mNG fluorescence of three replicates with light-incubated samples indicated by yellow points, and dark-incubated in black. **d,** Images of markerless Synpcc7942_0252-mNG tagged populations with and without 8 hours incubation in the dark **e,** Images of populations of Gap2-, Prk-, Cp12-mNG tagged *S. elongatus* markerless lines before and after one minute in darkness. **f,** Frames of time-lapse microscopy showing rapid relocalisation of mNG-tagged Prk protein upon the removal of light. **g,** Frames from time-lapse microscopy of the fusion of Prk-mNG-containing puncta. Arrows indicate puncta that coalesce within this timeframe. Scale bars: 1μm.

Cp12 is a known regulator of the Calvin cycle, thought to bind Gap2 and Prk to sequester them away from the cycle under conditions non-optimal for sugar production^31^. Tagging of Gap2 and Prk with mNG allowed us to visualise this process in real time, such that we could show the rapid relocalisation of these tagged proteins from a diffuse localisation pattern to distinct puncta within seconds of light being removed (Fig. 5b, Movies S1, S2 & S3). These puncta are highly dynamic, able to fuse, and appear to wet thylakoid membranes (Fig. 5c, Movie S4). These dynamic properties are widely associated with biomolecular condensates that form through liquid-liquid phase separation^32^, however further *in vivo* and *in vitro* supporting data is required to ascertain this^33^. Puncta still form in the absence of Synpcc7942_0252 (Figs. S2b & S2c) that we observed to interact with Gap2 and Prk, indicating that this interaction is not essential to the dark-induced protein relocalisation. This is in agreement with the low expression of Synpcc7942_0252 during the rapid relocalisation of Gap2 and Prk observed upon light to dark transition and would suggest that Synpcc7942_0252 has a role after Gap2/Prk puncta formation (Fig. 5c).

### Protein tagging provides further clues to protein function

By tagging previously uncharacterised proteins we stand to generate insights into where - and subsequently how - they function. For example, by tagging Synpcc7942_0011, we identified the brightest signals at the junctions between daughter cells as they separate from one another during the division process (Fig. 6a). This may indicate that Synpcc7942_0011 functions during the process of cell division. To investigate the function of Synpcc7942_0011 we used a modified CyanoTag plasmid vector to disrupt its genetic locus and create a knockout. Following the disruption of the *Synpcc7942_0011* locus (Fig. S2b) we did not observe significant changes in cell size or shape by fluorescence microscopy (Fig. 6a & 6b). Knockout lines were only slightly impaired in their growth rate (compared to wild-type and complemented strains), indicating that Synpcc7942_0011 is not required under our standard conditions (Fig. 6c). Further investigation is required to determine what role this previously uncharacterised protein is playing at sites of cell cleavage.

**Fig. 6:**
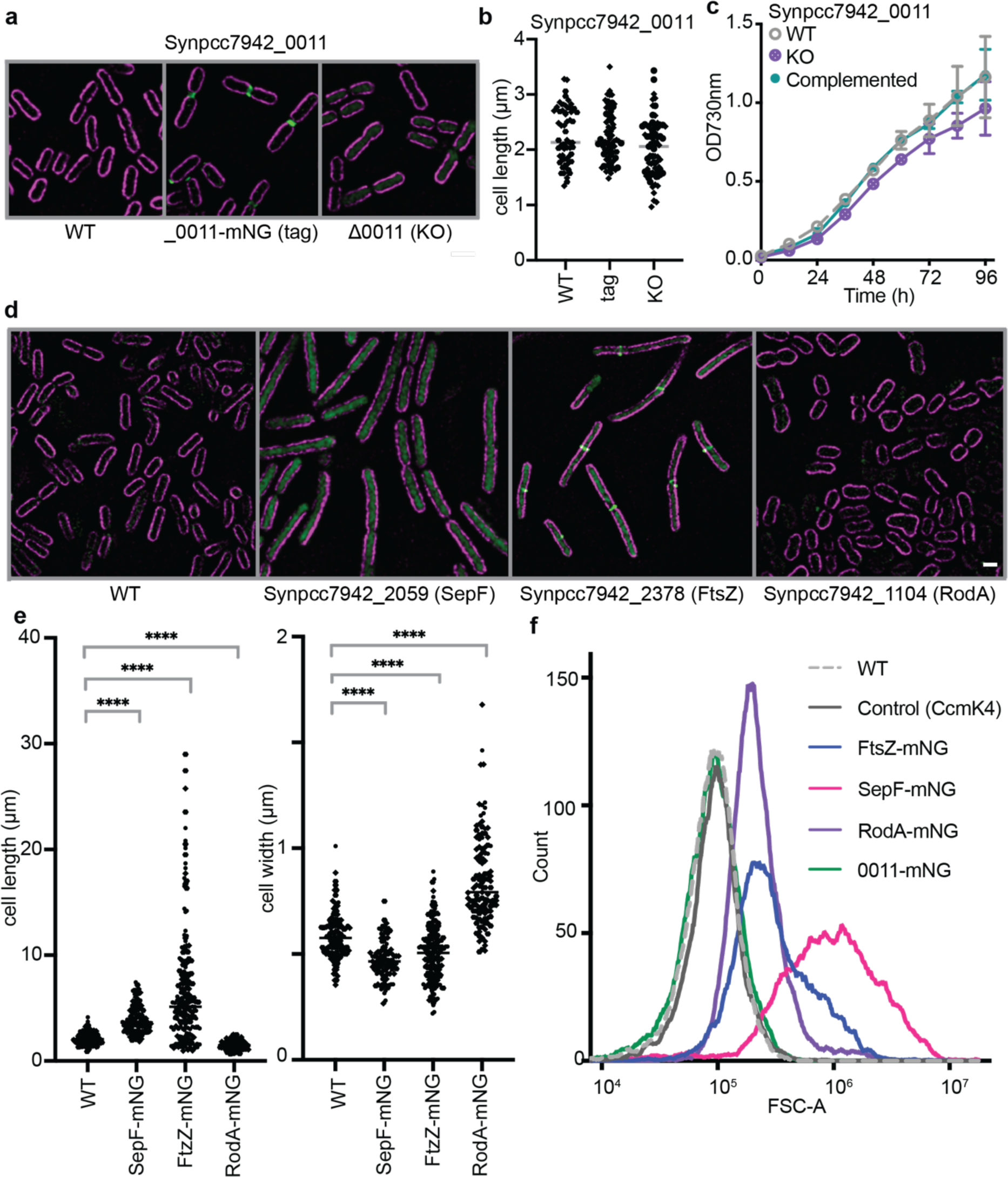
CyanoTag mutants provide insights into mediators of cell division and morphology. **a,** Fluorescence microscopy showing *S. elongatus* populations with tagged (middle) and disrupted *Synpcc7942_0011* loci (right). **b,** Analysis of cell dimensions of populations shown in a). Data points represent one of at least 25 cells measured from each group in each of two experiments. **c,** Growth of wild type, *Synpcc7942_0011* knockout and complemented lines measured by absorbance of populations at 730 nm. **d,** Fluorescence microscopy of markerless CyanoTag mutant targeting the *Synpcc7942_2059, Synpcc7942_2378 & Synpcc7942_1104* loci. Where Synpcc7942_2378 (FtsZ) is tagged, sites of mNG signal density at potential cleavage furrows, likely correspond to the z- ring structure. **e,** Analysis of cell dimensions of populations in e). At least 100 cells measured per condition across three experiments. **f,** Forward scatter (area: FSC-A) parameter of indicated cell populations measured by flow cytometry. All scale bars: 1μm.

In addition to localisation information, we observed several striking cell shape phenotypes during fluorescence imaging of three CyanoTag lines; tagging of Synpcc7942_2059 (SepF) or Synpcc7942_2378 (FtsZ) resulted in elongated cells whereas tagging Synpcc7942_1104 (RodA) resulted in shorter, wider cells (Fig. 6d, 6e & 6f). In bacterial cell division, the FtsZ protein forms a contractile ring at the midpoint of a growing cell, constricting to cause the separation of daughter cells. The SepF protein is thought to play a role in assembling FtsZ filaments and recruiting them to the cell membrane^34^. Genetic disruption of these two proteins has been shown previously to result in elongated cells similar to those we observe in the corresponding CyanoTag lines^35^. Synpcc7942_1104 is annotated as a functional homologue of RodA, a protein thought to be involved in cell wall synthesis and important for maintaining elongated cell shapes^36^. The morphological changes we observe in these would be consistent therefore with the proteins’ functions being disrupted by the addition of the C-terminal mNG tag. Fusion tags’ influence on protein function is a widely recognized caveat of fluorescent protein tagging^11^, however as demonstrated here this impairment does not always preclude the utility of these tagged lines, and may - in contrast - provide further insights into protein function.

## Discussion

### New insights and publicly available data and resources

We have developed the robust, high-throughput CyanoTag pipeline that can be used to generate *S. elongatus* cell lines expressing mNeonGreen-tagged proteins from their native loci. We demonstrated the use of these tagged lines to gain insight into a range of attributes for over 330 proteins, including their localisations, abundance and interactions. Amongst the insights we have gained from studying our CyanoTag lines were real-time observations of spatial and temporal regulation of two key Calvin cycle proteins, Prk and Gap2. Our AP-MS studies also identified a potential novel interactor of Prk and Gap2, though whether/how it regulates Calvin cycle dynamics will require further investigation. Additionally, the generated protein-protein interactome has robustly assigned proteins of unknown function to central, well-studied multimeric complexes, such as PSII and NDH-1. We were also able to identify a possible role in cell wall dynamics for a previously uncharacterised protein Synpcc7942_0011, initially clued via its striking localisation at the division sites of separating cells.

In this work, we report our initial pilot study targeting 400 of the 2,714 proteins encoded by the *S. elongatus* genome. With optimised high-throughput protocols, using a multiplexed format which has allowed significant reductions in resource and plastics usage, this pipeline is readily scalable. We are already expanding our libraries, working towards greater coverage of the proteome, offering more opportunities to observe novel biology and generate the first experimentally-defined whole cell interactome of a photosynthetic organism. Our data is available publicly via our online platform at https://morf-db.org/projects/York-Mackinder-Lab/MORF000032 which we will update as the project progresses and has the capability of integrating parallel datasets from compatible studies on *S. elongatus.* To further support the cyanobacterial research community, we are open to sharing our CyanoTag lines with other researchers, and have also created and shared detailed protocols for each step in our pipeline (included in Supplementary Material).

### Limitations of our datasets

As with all high-throughput strategies, there are some important limitations to be considered when working with our datasets and tagged lines. Scarless tagging of proteins at their endogenous loci has the great advantage of keeping native regulatory elements intact, but in any case, the addition of a 30 kDa tag at the C-terminus of a protein has the potential to impair normal protein function, localisation, interactions or expression. This may preclude homoplasmic modification (observed with RbcS, Table S1) or result in growth impairment (observed with SepF, Fig. 6). However, this did not prevent us from tagging at least one copy of our target gene in over 90% of attempted transformations (Fig. 1e) and observing correlation between our protein expression data and previously published estimates of protein copy number (Fig. 3e). In several cases, tagging interfering with protein function allowed us to make inferences about the target protein’s role, for instance an elongated cell shape implied a protein involved in cell separation had been disrupted (Fig. 6e & 6f).

Where markerless mutants cannot be generated or are too greatly impaired, a possible alternative is to use non-homoplasmic or marked CyanoTag lines maintained with kanamycin selection. In the course of developing the pipeline, we noted generally increased expression of mNG in marked mutants compared to their markerless counterparts (particularly when tagged protein expression was low; Fig. S2e), suggesting that either some expression of the second mNG-3xFLAG module (Fig. 1b) can occur or that the efficient *rrnB* terminator^37^ inserted downstream of the first mNG-3xFLAG consistently enhances gene expression/protein translation over the native terminator. Whilst this does not preclude the use of non-homoplasmic or marked mutants in further studies, it is an important caveat of their use, and is the reason our analyses in this manuscript focus on markerless lines.

The impact of the C-terminal tag, combined with the conditions in which we have imaged CyanoTag lines also play into the confidence that can be placed in the localisations we have ascribed. The images in our library were collected using the same set of growth and imaging conditions, and as such we are not necessarily observing protein localisation in the conditions where each protein is optimally expressed or at the location where it functions. The nutrient composition of the growth media affects a range of proteins, for instance the balance of plastocyanin and other cytochromes shifts based on copper availability^38^; hence our use of <300nM copper in our media may explain why we see low expression of plastocyanin (Table S1). Similarly, because we grew our lines in constant light, we observed low expression of the dark-induced Synpcc7942_0252 under our standard imaging conditions (Fig. 5d). Our observations of Gap2 and Prk also highlight how dynamic protein localisation can be; these proteins play their primary role in the Calvin cycle in the cytoplasm, but quickly relocalise to an alternate punctate location under our imaging conditions (Fig. 5e & 5f, Movies S1-4). We also acknowledge that, even with experimental replication, most of our images are a snapshot of one particular point during the log phase of each cell line’s growth and that our datasets are not capturing oscillations of protein expression that are known to occur even under constant growth conditions^39^. We encourage the use of our tagged lines in further studies of specific proteins or responses to conditions.

### Conclusion

Photosynthetic organisms play a central role in ecosystem stability and global biogeochemical cycling. There is a growing need to accelerate our understanding of photosynthetic microbes and to use this knowledge to address huge global challenges relating to food security, threatened ecosystems and the escalating climate crisis. The CyanoTag project aims to help unlock the potential that understanding cyanobacterial biology can offer to these vital areas.

## Methods

### Cyanobacterial culture conditions

Mutants were generated on a WT *Synechococcus elongatus* PCC 7942 background. Cultures were maintained at 30℃ under constant light conditions (50 µmol photons/m^2^/s) on 1.5% agar BG-11 media plates. Liquid cultures were maintained in 1 mL volumes of BG-11 in deep 96 well plates and shaken on an orbital plate shaker at 150 rpm.

### Plasmid design and Golden Gate cloning

Two CyanoTag template plasmids, pLM433 and pLM434, were constructed. Both plasmids contained two copies of the cytotoxin-encoding gene (*ccdB*) each flanked by *BspQ*I (pLM433) or *BsaI* (pLM434) restriction sites. Present between the *ccdB* sequences were a mNG/3xFLAG tagging module, a kanamycin selection marker (*nptI*) and a negative selection marker (*codA*) followed by a second copy of the mNG/3xFLAG module. To generate plasmids capable of integration at the 3’ end of each target gene, a Golden Gate cloning reaction was used to replace the *ccdB* sequences with ∼800 base pair homology arms that had been amplified from genomic DNA by PCR. Target genes were selected on the basis of them being (1) known or associated with photosynthesis (see ^40^ for a review), (2) candidate CO_2_-concentrating mechanism (CCM) and low-CO_2_ induced genes, including those identified from transcriptomic and proteomic studies^19,41,42^, or (3) compartment markers.

### Generation and selection of mutants

*S. elongatus* PCC7942 cultures were transformed by incubation with plasmids overnight at 30℃ in the dark. Selection on BG-11 agar plates containing 25 µg/mL kanamycin was used to isolate transformed cells which were then grown in wells of deep 96-well plates in 1 mL BG-11 medium containing 25 µg/mL kanamycin. Cyanobacteria were selected on increasing concentrations of kanamycin (50, 100 and finally 200 µg/mL) and homoplasmic marked mutants were identified using colony PCR. “Markerless” mutants were then generated by homologous recombination between the two mNG/3xFLAG module via selection on BG-11 agar containing 0.1g/L 5-FC. Homoplasmic markerless mutants were identified using colony PCR. *Synpcc7942_0011* & *Synpcc7942_0252* knockout lines were generated using similar methods; modified plasmids carrying a kanamycin resistance cassette flanked by homology arms corresponding to the target gene were transformed to induce insertional mutants (Fig. S2b). To complement the *Synpcc7942_0011* knockout, the Synpcc7942_0011 coding sequence was amplified from the *S. elongatus* PCC 7942 genome and inserted into the pSyn_6 vector which contains NS1 (neutral site 1) homologous recombination regions and a strong constitutive *psbA* gene promoter. Following transformation, colonies were selected with 50 μg/ml spectinomycin. Knockout and complemented lines were validated by colony PCR and sequencing.

### Fluorescence microscopy

Super-resolution imaging was performed using a Zeiss Elyra 7 microscope, using a lattice SIM^2^ method, with the ‘scale to raw image’ parameter enabled. Images were further processed using Fiji (Image J). Comparison of the mNG and cellular autofluorescence (mostly from chlorophyll) signal distribution was used to determine subcategories of protein localisation. Where mNG signal was low and the resultant mNG localisations looked similar to that of WT cells, no localisation descriptors were assigned. Fluorescence intensity profiles and quantifications of cell dimensions were performed using the measurement tools in Fiji. For time-lapse imaging, samples were incubated in the microscope chamber under 50 µmol photons/m^2^/s LED light until immediately prior to recording.

### Quantification of protein abundance

*S. elongatus* tagged mutant colonies were transferred to 96-well microplates containing BG-11 medium and incubated at 30℃ under 50 µmol photons/m^2^/s light for two days, after which they were analysed using a Cytoflex S cytometer. Live cells were gated based on their autofluorescence and the median fluorescence intensity of this live cell population in the FITC channel was used as a proxy for mNG fluorescence.

### Affinity purification-mass spectrometry

Lines were prioritised for AP-MS based on having significant protein expression, interesting localisations and/or being involved in diverse subcellular processes. Tagged lines were grown in duplicate to log phase before being pelleted (∼30 mg pellets) and snap-frozen in liquid nitrogen. Pellets were resuspended in an immunoprecipitation (IP) buffer (200 mM D-sorbitol, 50 mM HEPES, 50 mM KOAc, 2 mM Mg(OAc)_2_, 1 mM CaCl_2_) containing protease inhibitors (Roche cOmplete EDTA-free), 2% digitonin, 1 mM PMSF, 0.5 mM NaF and 0.15 mM Na_3_VO_4_. Cells were lysed by agitation with 0.4-0.6 mm glass beads for 6 seconds followed by a 10 second incubation on ice for a total of 15 minutes. Lysate was incubated with anti-mNeonGreen nanobody-Trap Agarose beads (ChromoTek) according to the manufacturer’s instructions, for 60 minutes on a rotating platform at 20 rpm. Beads were washed three times with IP buffer containing 0.1% digitonin followed by a final wash in the absence of digitonin. All incubation steps were carried out at 4℃. Purified proteins were digested on-bead with the addition of 100 ng of Promega sequencing grade trypsin (V5111) and incubation at 37°C overnight. Peptides were loaded onto EvoTip Pure tips for nanoUPLC using an EvoSep One system. A pre-set 100 SPD gradient was used with a 8 cm EvoSep C_18_ Performance column (4 cm x 150 μm x 1.5 μm). The nanoUPLC system was interfaced to a timsTOF HT mass spectrometer (Bruker) with a CaptiveSpray ionisation source (Source). Positive PASEF-DIA, nanoESI-MS and MS^2^ spectra were acquired using Compass HyStar software (version 6.2, Thermo). Instrument source settings were: capillary voltage, 1,500 V; dry gas, 3 l/min; dry temperature; 180°C. Spectra were acquired between *m/z* 100-1,700. DIA fragmentation of 25 Th windows were used covering *m/z* 400- 1201 and an ion mobility range of 0.6-1.6 1/k0. Collision energy was interpolated between 20 eV at 0.5 V.s/cm^2^ to 59 eV at 1.6 V.s/cm^2^. Resulting data in .d format were searched using DIA-NN software (version 1.8) against a an *in-silico* predicted spectral library generated from the *S. elongatus* PCC 7942 subset of UniProt. DIA-NN peptide identifications were compiled with KNIME and data filtered to 1% FDR. Two peptides (LATSPVLR & IAQVNLSR) likely corresponding to trypsin autolysis peptides were stripped from the results to prevent false positives. All mass spectrometry data sets, along with DIA-NN search in-puts and results files are referenced in ProteomeXchange (PXD049961) and are available to download from MassIVE (MSV000094128) [doi:10.25345/C5VQ2SN3R].

### AP-MS data analysis

DIA-NN-derived, non-normalised protein group quantification values from all samples were run through a CompPASS package in R Studio and a continuous measurement variation of SAINT analysis in Ubuntu. Interactions which fell in both the top 1% WD score (CompPASS) and 7% AvgP score (SAINT) were defined as high confidence interactors. These were inputted into Cytoscape for interactome generation.

### More detailed supplementary methods are supplied and will be updated and shared publicly

## Acknowledgements

We thank the University of York Technology Facility, particularly Karen Hogg for flow cytometry assistance and John Davey for initial bioinformatics support. We also thank Gavin Thomas and Joyce Bennet for support making our datasets publicly available via MORF. We are also grateful for Luning Liu’s advice at an early stage of the project. We would also like to acknowledge undergraduate students Curtis Bendelow, Ezra Porter, Philip Loesel, Riley Bell and Zak Forster who helped make some of the plasmids as part of their BSc projects, and masters student Huw Stephens who helped generate several of the markerless mutants. The authors also thank James Barrett for feedback that improved the manuscript. The York Centre of Excellence in Mass Spectrometry was created thanks to a major capital investment through Science City York, supported by Yorkshire Forward with funds from the Northern Way Initiative, and subsequent support from EPSRC (EP/K039660/1; EP/M028127/1).

## Author Contributions

The CyanoTag pipeline was initially conceived by LCMM, established by GZ, then expanded and optimised by AJP. The majority of the CyanoTag mutants were generated by GZ and AJP: GZ constructed around 300 plasmids and generated around 250 of the marked mutants and 150 of the markerless mutants used in this work; AJP constructed around 40 plasmids, 80 marked and 170 markerless lines; MD constructed around 40 plasmids. AJP carried out PCR-based validations of all 650+ CyanoTag lines. Western blotting by MD. All fluorescence microscopy included was carried out by AJP, with support from GC. All flow cytometry was carried out by AJP. Affinity purifications were performed by MD and subsequent mass spectrometry analysis by AAD. The resulting interactome was assembled from these data by MD. Knockouts of Synpcc7842_0011 & _0252 were generated by GZ. Growth analysis on Synpcc7842_0011 knockouts by GZ. The manuscript was written, data collated, and figures drawn by AJP, with contributions from MD & LCMM. VS developed the data-sharing interface for this project. This project was initiated via funding through a University of York pump priming award to LCMM and funded through a UKRI Future Leaders fellowship (MR/T020679/1) awarded to LCMM, who provided overall supervision of the work.

## Supplementary information

### Supplementary Figures

**Fig. S1.**
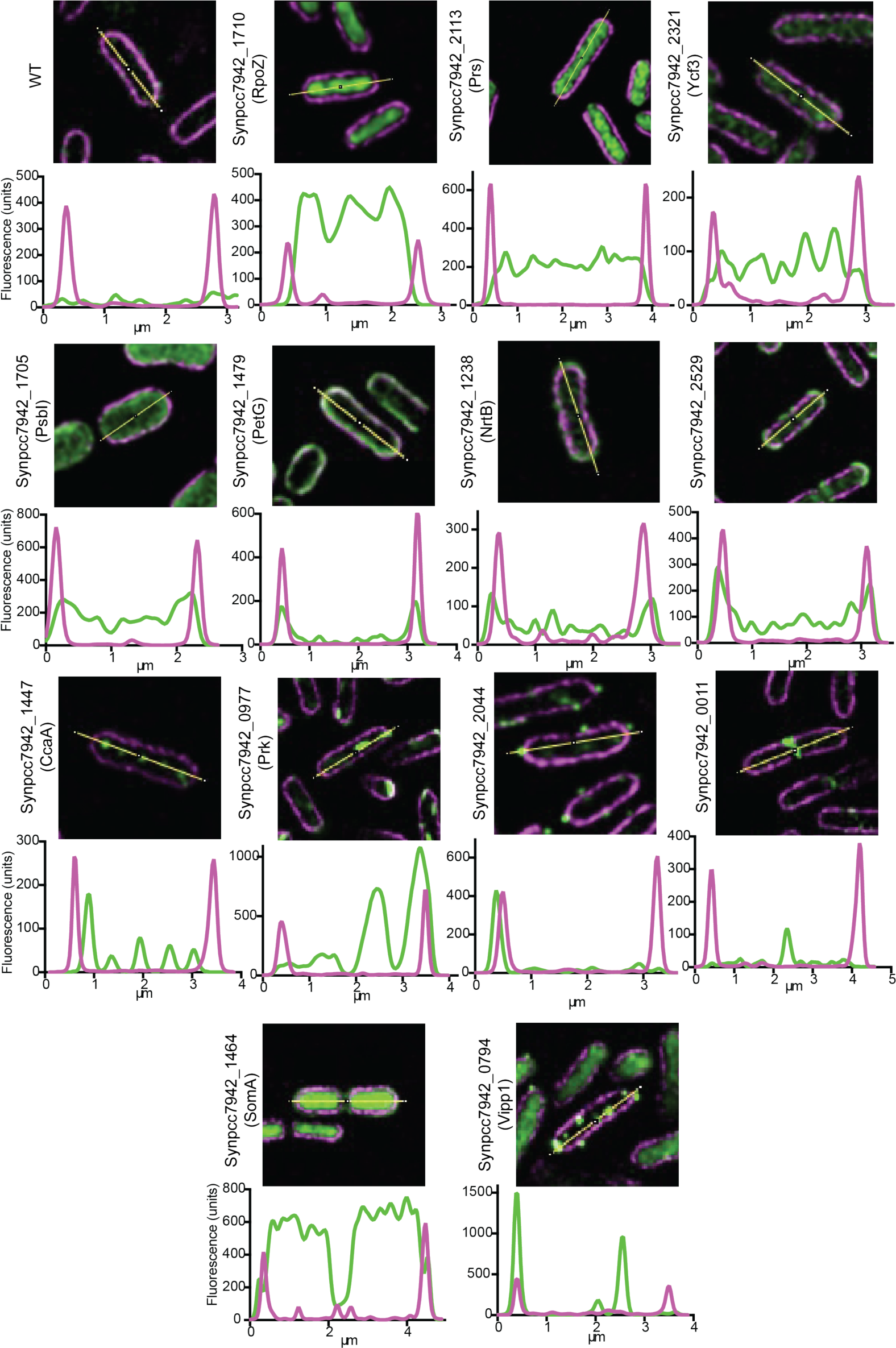
Example fluorescence microscopy images used in Fig. 2b & 3a of tagged proteins accompanied by the transects across which the intensity of the autofluorescence signals (magenta) and mNG fluorescence (green) were measured, Fluorescence intensity values (doubled for the mNG channel) across the transect are plotted below each image.

**Fig. S2:**
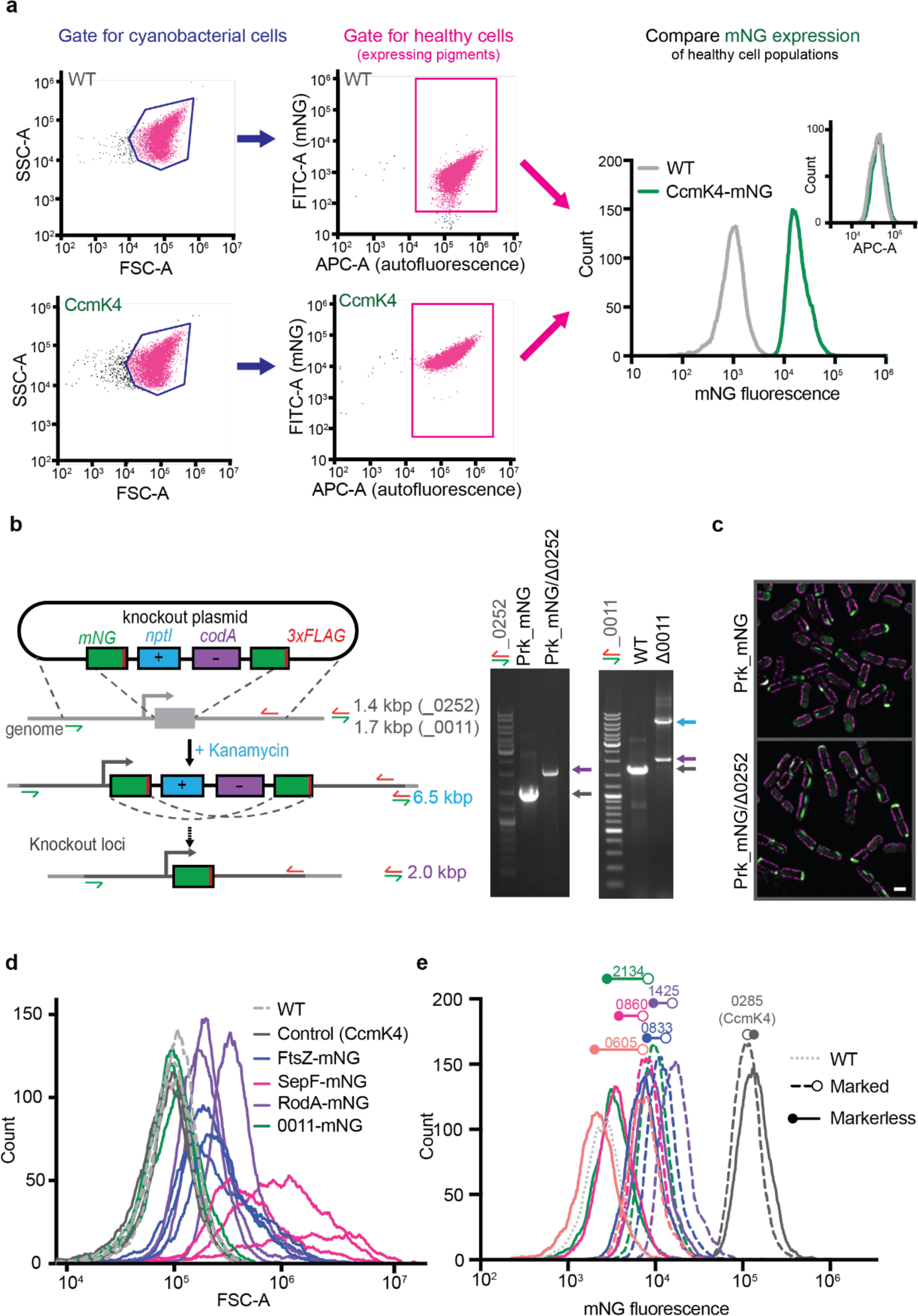
**a,** Example of flow cytometry gating approach used to quantify mNG expression in CyanoTag lines. APC-A: allophycocyanin (area parameter) **b,** Left: Disruption of *Synpcc7942_0252* & *Synpcc7942_0011* loci using modified CyanoTag vectors. Right: PCR validation of disruption of target loci. **c,** Fluorescence microscopy images of Prk puncta forming in the presence and absence of *Synpcc7942_0252.* Scale bar: 1μm **d,** Forward scatter (area: FSC-A) parameter of indicated cell populations described in Fig 6 measured by flow cytometry, showing three biological replicates. **e,** Flow cytometry-based quantitation of the mNG fluorescence of equivalent marked and markerless populations.

**Fig. S3:**
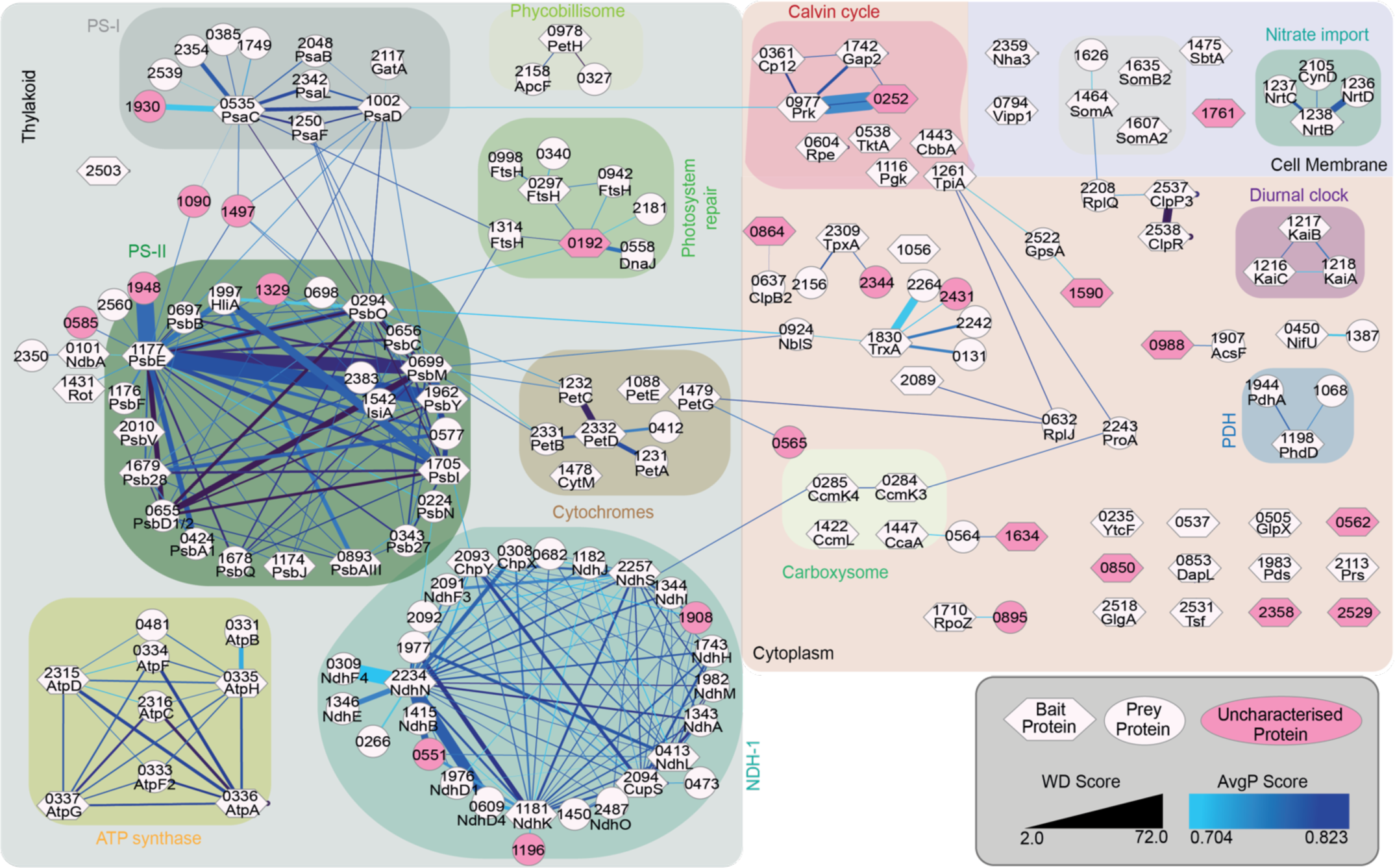
Preliminary protein interactome of *S. elongatus* based on data from 82 bait proteins. Labels on nodes denote proteins by their four number gene identifier and by an abbreviated name if one exists. Node shape corresponds to bait or prey status and edge width and colour represent WD and AvgP scores respectively. Known complexes are highlighted by background colour and currently uncharacterised protein nodes are shaded pink.

## Supplementary Movies

**Movie S1:** Time lapse fluorescence imaging of live markerless mNG-tagged Synpcc7942_0977-expressing (Prk-mNG) CyanoTag line upon withdrawal of light. Images were collected every 0.5 s. Magenta: cellular autofluorescence, green: mNG. Scale bar: 5μm

**Movie S2:** Time lapse fluorescence imaging of live markerless mNG-tagged Synpcc7942_1742-expressing (Gap2-mNG) CyanoTag line upon withdrawal of light. Images were collected every 0.5 s. Magenta: cellular autofluorescence, green: mNG. Scale bar: 5μm

**Movie S3:** Time lapse fluorescence imaging of live markerless CyanoTag lines expressing mNG-tagged Prk (Synpcc7942_0977) Gap2 (Synpcc7942_1742), and Cp12 (Synpcc7942_0361). Images were collected every 0.5 s for one minute following withdrawal of light. Top: merge of cellular autofluorescence (magenta) and mNG fluorescence (green). Bottom: mNG channel only. Scale bar: 5μm

**Movie S4:** Fusion of Prk-mNG puncta observed via time lapse fluorescence imaging of live markerless mNG-tagged Synpcc7942_0977-expressing (Prk-mNG) CyanoTag line upon withdrawal of light. Images were collected every 0.5 s. Magenta: cellular autofluorescence, green: mNG. Scale bar: 1μm

## Supplementary Tables

**Table S1:** Summary data for all CyanoTag lines used in this study.

**Table S2:** Summary of all interactions detected in affinity purification-mass spectrometry experiments

**Supplementary Methods** supplied as a PDF

