## Supplementary Methods for "CyanoTag: Discovery of protein function facilitated by high-throughput endogenous tagging in a photosynthetic prokaryote"

### CyanoTag Methods (v1.0) 26.02.2024

#### Step-by-step protocols for the generation and analysis of scarlessly tagged proteins in *Synechococcus elongatus* PCC7942

*Written by Dr Abi Perrin, based on CyanoTag protocols developed by Abi Perrin, Guoyan Zhao, Matt Dowson and Luke Mackinder.*


Background
Generating CyanoTag lines
- 1.1 Plasmid Cloning
  - 1.1.1 Backbone plasmid preparation
  - 1.1.2 Homology arm design and amplification
  - 1.1.3 Golden Gate Cloning
  - 1.1.4 E. coli transformation
  - 1.1.5 Plasmid validation and purification
- 1.2 Cyanobacterial strain generation
  - 1.2.1 S. elongatus transformation
  - 1.2.2 Marked mutant selection
  - 1.2.3 Marked mutant validation
  - 1.2.4 Markerless mutant selection
  - 1.2.5 Marked mutant validation
- 1.3 Storage and maintenance
  - 1.3.1 Maintenance on BG-11 agar
  - 1.3.2 Cryopreservation

Analysis of CyanoTag lines
- 2.1 Fluorescence microscopy
  - 2.1.1 Sample Preparation
  - 2.1.2 Fluorescence imaging
- 2.2 Flow Cytometry
- 2.3 Affinity Purification - Mass Spectrometry
  - 2.3.1 Preparing S. elongatus cell lysates
  - 2.3.2 Affinity Purification
  - 2.3.3 Mass Spectrometry

Notes
- Recommended equipment and materials
  - Equipment details
  - Reagents
- Reducing single-use plastic waste
  - Plastics Washing protocols
  - 48-well agar plate divider usage

Appendices
- A1: Media and Buffer Recipes

BG-11 recipes
Affinity purification buffers
LB24
A2: Plasmid Sequences and Maps
pLM433
pLM434
A3: Macro for making montages from single frames of images from the Elyra 7 in Fiji

### Background

Here we describe a high-throughput approach for scarless endogenous fluorescent protein tagging in the model cyanobacterium *Synechococcus elongatus* PCC7942. To date we have used this platform to fluorescently tag over 500 *S. elongatus* proteins and have used the resulting cyanobacterial cell lines to determine proteins' subcellular localisation, track relative protein abundances and to elucidate protein-protein interaction networks. Most steps can be multiplexed and are amenable to 96-well formats.

Our data have already provided novel insights into a diverse range of processes relevant to cyanobacteria and to more broadly to photosynthetic and/or bacterial life. We hope these lines, the insights they provide and the optimised methods we have developed in the course of this work will be a valuable resource to cyanobacterial cell biologists, and more widely!

### Generating CyanoTag lines

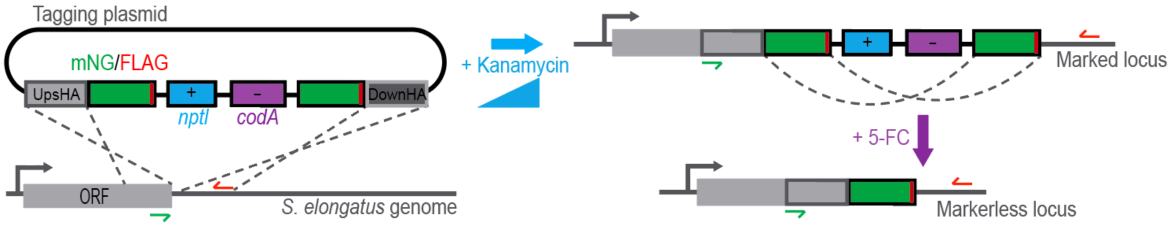


*Fig 1a: Genetics underlying the CyanoTag modification pipeline*

#### 1.1 Plasmid Cloning

The construction of CyanoTag plasmids involves the insertion of two homology arms (used to target the construct to the target genetic locus) into one of two CyanoTag vectors (pLM433 or 434) using a Golden Gate methods with the restriction enzymes BspQI or BsaI.

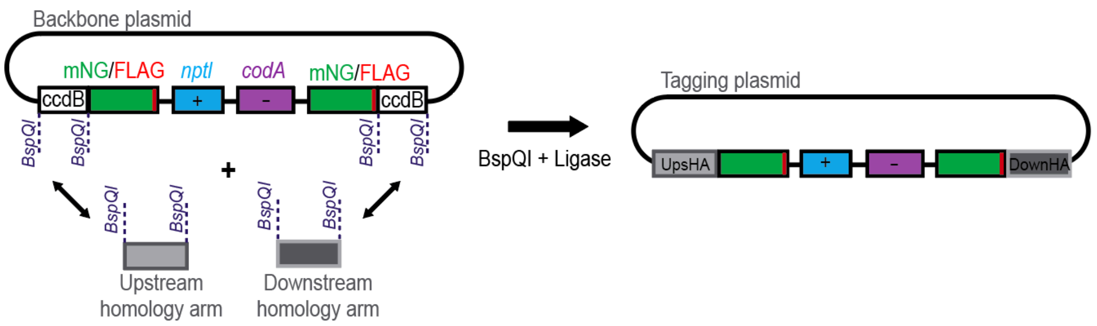


*Fig 1.1a: Golden Gate Cloning of CyanoTag Vectors. N.B. A BsaI mediated Golden Gate reaction is used where BspQI sites in homology arms would preclude cloning via this approach.*

#### 1.1.1 Backbone plasmid preparation

*Backbone plasmids maps and sequences are provided below and at the time of writing are in the process of being deposited are in Addgene*

1. Transform the backbone plasmid (pLM433 or 434) into an *E.coli* strain that is resistant to toxicity from the ccdB gene product (e.g. strains that contain the F plasmid and strains otherwise engineered to be resistant - see here for a list)
2. Grow transformed bacteria (from a colony or glycerol stock) in liquid culture. We use LB24 medium with 25 µg/µL carbenicillin (or ampicillin) and 10 µg/µL kanamycin
3. Purify plasmids (e.g. by miniprep), aiming for a final DNA concentration of at least 200 ng/µL.

#### 1.1.2 Homology arm design and amplification

*N.B. You may prefer to replace these amplification steps by ordering synthetic homology regions to clone directly into the CyanoTag backbone vectors. We have routinely used homology arms of ~400 bp without noticeable effects on transformation efficiency.*

##### 1.1.2.1 Primer design

*We have already designed primers and cloning approaches for most genes. Please contact the Mackinder lab if you would like us to share these with you.*

1. Search the regions up to ~800 bp before and after the stop codon of your target gene for the presence of BspQI (***GCTCTTC***) or BsaI (***GGTCTC***) restriction enzyme recognition sites (check in both orientations)
2. Based on your results, choose a cloning method based on BspQI and pLM433 or BsaI and pLM434; BspQI sites in your homology regions are likely to preclude cloning using BspQI/pLM433, BsaI sites in your homology regions are likely to preclude cloning using BsaI/pLM434. If there are both BspQI and BsaI sites close to the stop codon, you can synthesise the homology arm with a single base pair change to remove the restriction site.
3. For the upstream HA, choose ~20 base regions for:
    - forward primer to anneal ~400-800 bp before the stop codon
    - reverse primer directly before the stop codon

    For the downstream HA, choose ~20 base regions for:

    - forward primer to anneal directly after the stop codon (the stop code is not included)
    - reverse primer starting ~400-800 bp after the stop codon
4. Append the following adapter sequences to each of the four primers before ordering

*Table 1.1.2.1a: Adapter sequences for CyanoTag primers*

| **Cloning method** | **Upstream HAs** | | **Downstream HAs** | |
|---|---|---|---|---|
| | **Forward adapter 5'-3'** | **Reverse adapter 5'-3'** | **Forward adapter 5'-3'** | **Reverse adapter 5'-3'** |
| **BspQI/pLM433** | TATA**GCTCTTC**AACT | TATA**GCTCTTC**AGCC | TATA**GCTCTTC**ATAG | TATA**GCTCTTC**AAAG |
| **BsaI/pLM434** | TATA**GGTCTC**AGACT | TATA**GGTCTC**ACGCC | TATA**GGTCTC**AGTAG | TATA**GGTCTC**ACAAG |

##### 1.1.2.2 gDNA preparation

1. Grow a culture of wild-type *S. elongatus*. We use 50mL of culture with an $OD_{730}$~1.
2. Pellet cells by centrifugation at 1500x g for 15 minutes. Pellets can be used immediately or snap frozen
3. Extract genomic DNA. We use the Promega Wizard™ Genomic DNA Purification Kit, aiming for a final DNA concentration of at least 50 ng/µL.
4. Store at 4℃. Ideally use this within a couple of months.

##### 1.1.2.3 PCR amplification of HAs

1. Amplify homology arms (upstream and downstream) from gDNA using a high-fidelity, proofreading polymerase. We use Phusion and set up the PCR reaction and thermocycler as follows:

*Table 1.1.2.3a: PCR reagents for homology arm amplification (Phusion)*

| **Reagent** | **Per reaction (µL)** | **x100 (µL)** (for a 96 well plate) |
|---|---|---|
| 5× Phusion HF Buffer | 10 | 1000 |
| dNTPs (10 mM) | 1 | 100 |
| gDNA (~50 ng/µL) 1 | 1 | 100 |
| DMSO (100%) | 1 | 100 |
| Phusion DNA Polymerase | 0.5 | 50 |
| Water (molecular grade) | 31.5 | 3150 |
| F primer (10 µM) | 2.5 | Do not include in master mix |
| Reverse primer (10 µM) | 2.5 | Do not include in master mix |

*Table 1.1.2.3b: Thermocycler program for homology arm amplification PCR (Phusion)*

| **Temperature** | **Time (s)** | **Cycles** |
|---|---|---|
| 98°C | 60 s | 1 |

| | | |
|---|---|---|
| 98°C | 10 s | x32 |
| 55°C | 30 s | |
| 72°C | 30 s | |
| 72°C | 10 min | 1 |

2. Check the success of the amplification by analysing 5 µL of the reaction by gel electrophoresis.
   If there is a single band of the expected size, purify the remaining DNA using a PCR cleanup kit (details in Plasmid cloning reagents) and measure the concentration of each HA

##### 1.1.3 Golden Gate Cloning

1. Combine backbone plasmid and homology arms in single Golden Gate reaction

*Table 1.1.3a: Golden Gate cloning mixture*

| **Reagent** | **Per reaction (µL)** | **x100 (µL)** (for a 96 well plate) |
|---|---|---|
| 10× T4 DNA ligase buffer | 0.5 | 50 |
| Upstream HA (~100 ng/µL) | 0.9 | - |
| Downstream HA (~100 ng/µL) | 0.9 | - |
| Backbone vector pLM433* (~200 ng/µL) | 2 | 200 |
| BspQI* (10 U/µL) | 0.25 | 25 |
| T4 Ligase | 0.125 | 12.5 |
| Water (molecular grade) | 0.325 | 32.5 |

**For assembly with pLM434 as the backbone vector, use BsaI instead of BspQI.*

*Table 1.1.3b: Thermocycler program for Golden Gate reactions*

| **Temperature** | **Time (s)** | **Cycles** |
|---|---|---|
| 37°C | 15 min | 1 |
| 37°C | 5 min | x20 |
| 16°C | 5 min | |
| 37°C | 5 min | 1 |
| 65°C | 25 min | 1 |

#### 1.1.4 *E. coli* transformation

1. Add 15 - 30 µL chemically competent *E. coli* (N.B. this must be a strain that is ***not*** resistant to ccdB) to each completed Golden Gate reaction and incubate the mixture on ice for 30 minutes
2. Heat-shock the plate/tubes for 90 seconds at 42°C in a thermocycler.
3. Incubate the cells on ice for another 2 minutes then transfer them to new vessels (deep well plates or microcentrifuge tubes) containing 250 µL SOC buffer per transformation. Shake the cells for 1 hour at 37°C
4. Plate 150 µL onto LB agar plate containing 25 µg/mL ampicillin and 10 µg/mL kanamycin and incubate the plate overnight at 37°C

#### 1.1.5 Plasmid validation and purification

1. Check colonies by PCR using primers
   a. oLM617: ACAAAGATCACGACATCGACTAT
   b. oLM618: CCGCTGCCACCCAGATCG

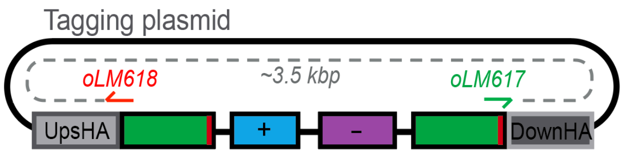


*Fig 1.1.5a: Check PCR used to determine successful integration of homology arms in Golden Gate cloning.*

We use a Taq-based master mix for this as follows.

*Table 1.1.5a: Colony PCR reagents (Vazyme)*

| **Reagent** | **Per reaction (µL)** | **x100 (µL)** (for a 96 well plate) |
|---|---|---|
| Primer oLM617 (10 µM) | 0.5 | 50 |
| Primer oLM618 (10 µM) | 0.5 | 50 |
| 2× Vazyme master mix | 5 | 500 |
| Water (molecular grade) | 4 | 400 |
| *E.coli* colony | n/a | Do not include in master mix |

*Table 1.1.5b: Thermocycler program for colony PCR (Vazyme)*

| **Temperature** | **Time (s)** | **Cycles** |
|---|---|---|
| 95°C | 10 min | 1 |
| 94°C | 60 s | x32 |
| 55°C | 30 s | |
| 72°C | 3 min | |
| 72°C | 10 min | 1 |

2. Check the products using gel electrophoresis - bands should be around or above 3kb in size.
3. Grow validated colonies overnight in 1-5 mL (depending on sample numbers and capacity) LB24 with 25 µg/mL ampicillin and 10 µg/mL kanamycin in a 37°C shaker.
4. Extract the plasmid DNA from the cells by using a miniprep kit (details in Plasmid cloning reagents), aiming for a final concentration of >50 ng/µL.
5. Following purification, check successful integration of homology arms by DNA sequencing. We use the following primers
    a. oLM352: CGACACGGAAATGTTGAA
    b. oLM349: GCTTGGAGCGAACGACC

#### 1.2 Cyanobacterial strain generation

##### 1.2.1 *S. elongatus* transformation

1. Grow a culture of wild type *S. elongatus* up to an $OD_{730nm}$ of ~0.5-1.0. We culture these cyanobacteria in BG-11 medium (see below) in glass flasks, on a shaking platform (150 rpm) in an incubator at 30°C under continuous illumination of 50 µmol photons/m$^2$/s and use 600 mL culture to transform a 96 well plate.
2. Harvest the cells by centrifugation at 1500 x*g* for 15 min.
3. Wash the cells by resuspending the pellet in 50 mL of 10 mM NaCl, then pelleting the cells by centrifugation at 1500 x*g* for 15 min.
4. Resuspend the pellet in BG-11 media (10 mL if transforming 96 wells) and transfer 100 µL of the cell suspension into each well of a microplate.
5. Add 10 µL of purified plasmid DNA (500-1000 ng) to each well and mix gently.
6. Incubate the cells at 30°C in the dark for 20-24 hours.
7. Plate at least 20 µL of the cells on BG-11 agar with 25 µg/mL kanamycin and incubate plates at 30°C under 50 µmol photons/m$^2$/s. Colonies should start appearing after about a week.

##### 1.2.2 Marked mutant selection

1. Transfer a single colony from each transformation into 1 mL BG-11 containing 50 µg/mL kanamycin. We use deep-well 96-well plates for this. Shake the plate (~300 rpm) at 30°C under 50 µmol photons/m$^2$/s for 2-3 days.
2. Increase the kanamycin selection: Transfer 100 µL of these cultures into 1 mL BG-11 containing 100 µg/mL kanamycin. Shake the plate (~300 rpm) at 30°C under 50 µmol photons/m$^2$/s for 2-3 days.
3. Increase the kanamycin selection again by transferring 100 µL of these cultures into 1mL BG-11 containing 200 µg/mL kanamycin. Shake the plate (~300 rpm) at 30°C under 50 µmol photons/m$^2$/s for 2-3 days.
4. Plate 5 µL of the cells on BG-11 agar with 200 µg/mL kanamycin and incubate plates at 30°C under 50 µmol photons/m$^2$/s. Colonies should start appearing after 2-3 days.

##### 1.2.3 Marked mutant validation

1. Check the target locus of each colony using a colony PCR

*Table 1.2.3a: S.elongatus Colony PCR reagents (Vazyme)*

| **Reagent** | **Per reaction (µL)** | **x100 (µL)** (for a 96 well plate) |
|---|---|---|
| Upstream HA Forward primer (10 µM) | 0.5 | Do not include in master mix |
| Downstream HA Reverse primer (10 µM) | 0.5 | Do not include in master mix |
| 2× Vazyme master mix | 5 | 500 |
| Water (molecular grade) | 4 | 400 |
| *S.elongatus* colony | n/a | Do not include in master mix |

*Table 1.2.3b: Thermocycler program for S.elongatus colony PCR (Vazyme)*

| **Temperature** | **Time (s)** | **Cycles** |
|---|---|---|
| 95°C | 10 min | 1 |
| 94°C | 60 s | x32 |
| 55°C | 30 s | |
| 72°C | 3 min | |
| 72°C | 10 min | 1 |

2. Check the products using gel electrophoresis. Marked bands should be around ~7 kb but we often see a degree of spontaneous marker removal and the resultant presence of a ~2.5kb band for the markerless locus as well as (or instead of) the ~7kb one.  Smaller bands - in our setup around 1.5kb - indicate the presence of the wild type locus in at least one copy of the gene.  The presence of any wild type locus will likely preclude subsequent marker removal.
3. Maintain validated marked mutants on BG-11 agar with 200 µg/mL kanamycin

#### 1.2.4 Markerless mutant selection

1. Transfer a single colony of a validated marked mutant to 1 mL BG-11 media (no antibiotics) and  shake the plate (~300 rpm) at 30°C under 50 µmol photons/m$^2$/s for 2 days until the cultures are pale green.
2. Plate 10-25 µl of each culture onto BG-11 agar containing 100 µg/mL 5-fluorocytosine (5-FC) and incubate plates at 30°C under 50 µmol photons/m$^2$/s. Colonies should start appearing after about a week.

#### 1.2.5 Marked mutant validation

1. Check the target locus of each colony using a colony PCR as detailed in 1.2.3 Marked mutant validation
2. Maintain validated marked mutants on BG-11 agar without antibiotics.

#### 1.3 Storage and maintenance

##### 1.3.1 Maintenance on BG-11 agar

1. CyanoTag mutants are usually viable stored at ambient temperature/light levels on BG-11 agar for a period of months.
2. We array our lines in 96-well plate format and replicate our libraries monthly using a Singer Rotor instrument.

##### 1.3.2 Cryopreservation

1. To freeze a 1 mL culture for longer term storage:
    a. Pellet cells by centrifugation at 5000 x*g* for 2 minutes
    b. Remove the supernatant
    c. Resuspend cells in 50-200 µL BG-11 + 5% DMSO
    d. Incubate cells on ice for 15 minutes before transferring to -70°C
2. To recover frozen isolates:
    a. Incubate cells on ice for 10-20 minutes to allow them to thaw
    b. For small numbers of lines: plate 10-50 µL cells directly onto BG-11 agar plates (with 200 µg/mL kanamycin for marked mutants) and incubate plates at 30°C under 50 µmol photons/$m^2$/s. Colonies should start appearing within a week.
    c. When thawing 96-well plates, transfer 15 µL into wells containing 1mL BG-11 (with 200 µg/mL kanamycin for marked mutants) and shake plate(s) at 30°C under 50 µmol photons/$m^2$/s. Wells should look green within 3-7 days.

### Analysis of CyanoTag lines

#### 2.1 Fluorescence microscopy

##### 2.1.1 Sample Preparation

1. Transfer colonies into 1mL BG-11 media and shake the plate (~300 rpm) at 30°C under 50 µmol photons/$m^2$/s for 48 hours
2. Prior to imaging, coat the wells of an imaging plate with poly-L-lysine. You can do this by incubating the cells with 0.01% (w/v) pol-L-lysine for 5 minutes, then removing the liquid and leaving the plate to dry overnight.
3. Transfer ~50 µL culture into each coated well and centrifuge plate at 3000 x*g* for 2 minutes. You can then leave the cells to settle for 30 minutes.
4. Cover the cells with ~150 µL 1.5% low melting point agarose (prepare in BG-11 medium) and leave to set.

##### 2.1.2 Fluorescence imaging

###### 2.1.2.1 Lattice SIM

We use a Lattice SIM method on a Zeiss Elyra 7 microscope to image our lines, illuminating the samples with the 488 nm laser and dividing the output signal between two cameras to capture the mNG signal and the cellular autofluorescence.

1. Ensure cameras are aligned and adjust them if needed. You can use the sample to do this (as opposed to beads).
2. Collect z stacks of desired fields of view.
3. Process using 3D $SIM^2$ algorithm, using “standard live” settings for the autofluorescence and “weak live” for the mNG channel. Use a test image to set an alignment matrix to be integrated into the processing. Ensure ‘scale to raw image’ is selected.
4. Alignment is often still not perfect for every image and using the channel alignment tool for your final image can help correct this.

###### 2.1.2.2 Image processing in Fiji

1. Select the desired frame of the z stack (where the cellular autofluorescence is clearest).
2. Adjust the brightness of the mNG channel to enable comparison between images. For the images in MORF I have set the default LUT range for this channel as 10-150, though have increased the upper threshold in images where the signal would be greatly oversaturated otherwise (i.e. high expressing lines).
3. We wrote a simple macro to process batches of these images into montages of each channel and a merge - see A2: Macro for making montages from single frames of images from the Elyra 7 in Fiji

#### 2.2 Flow Cytometry

1. Transfer colonies into 100 µL BG-11 media in a transparent microtitre plate. Cover the plate with a breathable seal and shake at 30°C under 50 µmol photons/$m^2$/s for 48 hours.
2. We used a Cytoflex S flow cytometer to analyse mNG fluorescence in these populations, running a well containing cleaning solution and another containing BG11 between each sample. Live cells were gated based on their autofluorescence and the median fluorescence intensity of this live cell population in the FITC channel was used as a proxy for mNG fluorescence

#### 2.3 Affinity Purification - Mass Spectrometry

##### 2.3.1 Preparing *S. elongatus* cell lysates

###### 2.3.1.1 Prepare cell pellets

1. Grow 50 mL of the CyanoTag line to an $OD_{730nm}$ of 0.5 (~ 5 days at 30°C under 50 µmol photons/$m^2$/s).

2. Harvest the cells by centrifugation at 4°C at 1500 x*g* for 15 min and completely remove supernatant. Aim for ~30 mg pellets.
3. Flash freeze the pellet in liquid $N_2$ for 90 seconds. Store the cells at -70°C until needed.

###### 2.3.1.2 Lyse cell pellets

*Perform the following steps at 4°C.*

1. Prepare and chill 2 mL microcentrifuge tubes containing ~170 mg glass beads 200 μL of AP buffer + PIs + 2% digitonin.
2. Transfer cell pellets into these chilled prepared tubes using a clean spatula (sterilise by washing in bleach followed by $H_2O$ between samples).
3. Vortex cells for 6 s, and keep on ice for 10 s; repeat the vortex and cool down process for 15 min. Ideally do this in a 4°C room.
4. Clarify lysate by centrifugation for 30 minutes at full-speed in a table-top centrifuge at 4°C.

##### 2.3.2 Affinity Purification

###### 2.3.2.1 Prepare NanoTrap Reagent

1. Resuspend mNeonGreen Nano-Trap beads by pipetting. Transfer 25 μL per purification to a 2 mL tube.
2. Place on a MagnaRack and remove storage liquid. Wash Nano-Traps with 0.5 mL ice-cold AP buffer + PIs (without digitonin).
3. Place tubes on MagnaRack and remove supernatant just before adding lysate.

###### 2.3.2.2 Capture proteins on beads

*Perform the following steps at 4°C.*

1. Transfer 200 μL of lysate to Nano-Traps being careful not to disturb the pellet or glass beads. Incubate for 1 hour on a rotating platform at 4°C at 20 rpm.
2. Place tubes on the MagnaRack and remove the supernatant (keep a sample of this for immunoblotting).
3. Wash the Nano-traps by adding 0.65 mL AP buffer + PIs and 0.1% digitonin, incubating for 3 minutes at 25 rpm on a rotating platform, then place tubes on the MagnaRack and remove the supernatant. Do this wash step 3 times.
4. Perform a final wash with 0.7 mL AP buffer + PIs (no digitonin) and remove all supernatant (keep a sample for immunoblotting)
5. These loaded beads can be stored at -20°C. Prior to on-bead digestion and mass spectrometry.

###### 2.3.2.3 On-bead digestion

1. Prior to mass spectrometry, add 100 ng of sequencing grade trypsin to each sample
2. Incubate samples overnight at 37°C overnight

##### 2.3.3 Mass Spectrometry

###### 2.3.3.1 Data acquisition

1. Samples were run through an 8 cm Performance column and analysed using parallel accumulation-serial fragmentation data independent acquisition (PASEF-DIA) with DIA fragmentation of 5 m/z windows covering 400-1201 m/z on a Bruker trapped ion mobility spectrometry time of flight (TimsTOF) mass spectrometer with a long gradient methodology.
2. Resulting data were converted to mzML using MSconvert, before searching using DIA-NN software with the *S. elongatus* PCC 7942 subset of UniProt and compiled with KNIME. These data were then filtered to 1% FDR.Two peptides (LATSPVLR & IAQVNLSR) likely corresponding to trypsin autolysis peptides were stripped from the results to prevent false positives.

###### 2.3.3.2 Data processing

1. We filtered the data by running non-normalised protein group quantification values using:
    a. a CompPASS package in R Studio, retaining those that fell within the top 1% in terms of their WD score.
    b. a continuous measurement variation of SAINT analysis in Ubuntu, retaining those that fell within the top 7% in terms of their AvgP score.
2. Interactions passing both thresholds were considered high-confidence and were used to generate an interactome in Cytoscape.

---

### Notes

#### Recommended equipment and materials

##### Equipment details

- ROTOR HDA instrument used for library maintenance, replication and arraying - Singer Instruments, used in combination with
    - PlusPlates - Singer Instruments (PLU-003)*
    - RePads 96 Long - Singer Instruments (REP-001)*
- Elyra 7 Super Resolution Microscope - Zeiss, used for live imaging in combination with
    - μ-Plate 96 Well Square Glass Bottom Imaging plate - Ibidi (89627)
- CytoFLEX S Flow Cytometer - Beckman Coulter
- timsTOF HT Mass Spectrometer - Bruker, used in combination with
    - nanoUPLC using an EvoSep One system
    - CaptiveSpray ionisation source

- Axygen® 96-well Clear Round Bottom 2 mL Polypropylene Deep Wells - Corning (P-DW-20-C)*
- Nunc™ Square BioAssay Dishes - ThermoFisher (240845)* *used with bespoke dividers to create 48-well agar plates see 48-well agar plate divider usage.*
- MagnaRack™ Magnetic Separation Rack - ThermoFisher (CS15000)
- Breathable Plate Sealing Film (Sterile) - Starlab (E2796-3015)
- Polyester Plate Sealing Film (Sterile) - Starlab (E2796-0714)

**Indicates that this product can be washed and reused (see Reducing single-use plastic waste for details and instructions)*

#### Reagents

##### Plasmid cloning reagents

- Wizard® Genomic DNA Purification Kit - Promega (A1120)
- Phusion™ High-Fidelity DNA Polymerase - ThermoFisher (F530)
- Wizard® SV 96 PCR Clean-Up System - Promega (A9340)
- QIAquick PCR purification kit - Qiagen (28104)
- BspQI - NEB (R0712)
- BsaI-HF®v2 - NEB (R3733)
- T4 DNA Ligase (thermostable) - NEB (M0202)
- 2 × Taq Master Mix (Dye Plus) - Vazyme (P112) for colony PCR checks
- QIAprep spin miniprep kit - Qiagen (27104)
- Wizard® SV 96 Plasmid DNA Purification Kit - Promega (A2250)

##### *S. elongatus* culture and selection reagents

- 5-Fluorocytosine (5FC) - Alfa Aesar L16496.MD
- Nystatin - Sigma (N6261)

##### 5-FC Preparation

Make a 10 g/L stock solution in DMSO. Aliquot and store at -20°C. Working concentration is 100 μg/mL (1 in 100 of the stock solution).

##### Imaging reagents

- Poly-L-lysine 0.1% (w/v) - Sigma (P8920) used 1 in 10
- UltraPure™ Low Melting Point Agarose - ThermoFisher (16520)

##### Affinity Purification & Mass Spectrometry reagents

- Digitonin - Sigma (D141)
- Glass beads, acid washed - Sigma (G8772)
- cOmplete Mini, EDTA-free protease inhibitor tablets - Roche (11836170001)
- mNeonGreen-Trap Agarose beads - ChromoTek nta-200
- Sequencing grade trypsin - Promega V5111

##### Digitonin Preparation

Prepare 10% digitonin solution by dissolving 100 mg Digitonin in 1 mL of dd$H_2O$ by adding $H_2O$ slowly to a large surface area of digitonin (spread equally across the length of a 2 mL tube). Heat at 60°C for 10 minutes if further dissolution required. Avoid generating bubbles with pipetting.

### Reducing single-use plastic waste

By multiplexing our protocols we are reducing resource usage per line, but there are still significant cost and new plastics savings to be made through reuse of ‘consumable’ items in the pipelines. Some examples of the approaches we have taken are below.

#### Plastics Washing protocols

1. Remove and decontaminate any culture or agar from the plate/surface
2. Incubate plates in 1% Virkon solution overnight (other disinfectants may also be appropriate)
3. Wash plates in water (standard tap) 3 times and incubate overnight in DI water
4. Dry the plates completely
5. Sterilise before use
    a. For polypropylene deep well plates, plate dividers and ROTOR Pads, autoclave
    b. For polystyrene plates (bioassay dishes and ROTOR plates), use UV light.

#### 48-well agar plate divider usage

We can significantly increase convenience and reduce cross-contamination and waste by dividing large agar plates into separate wells when selecting colonies on solid media. The workshop in the University of York Biology department made us some plate dividers using offcuts of polypropylene (which can be washed, autoclaved and reused). You can see how we use them below.

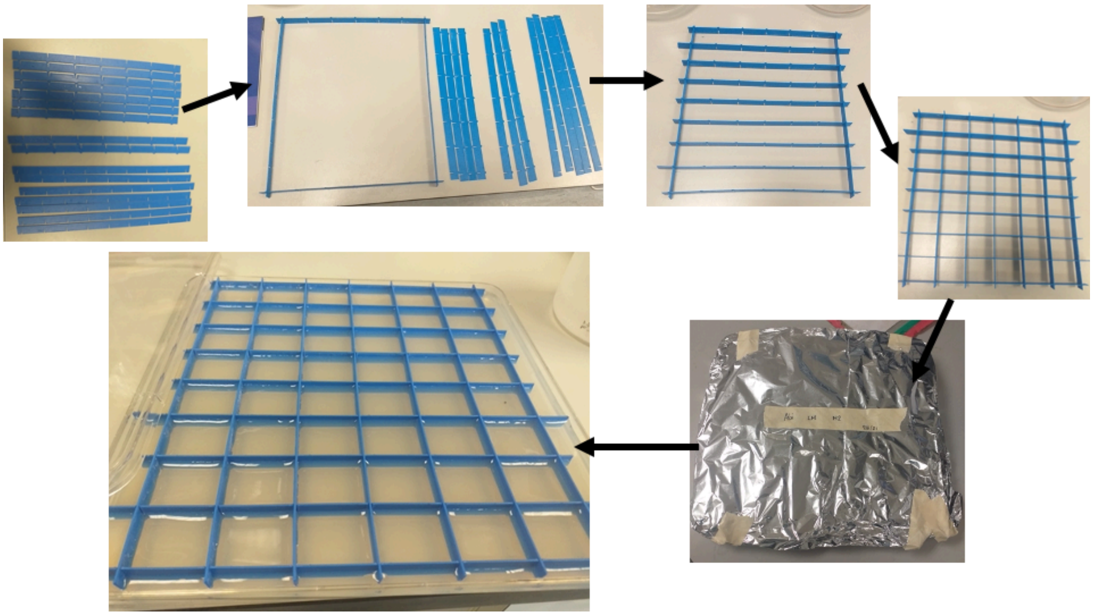

*Reusable plate dividers for CyanoTag pipeline*

---

### Appendices

#### A1: Media and Buffer Recipes

##### BG-11 recipes

#### 100x BG-11

| **Reagent** | **Quantity for 1 L of 100x BG-11** |
|---|---|
| $NaNO_3$ | 150 g |
| $MgSO_4.7H_20$ | 7.5 g |
| $CaCl_2$ | 2.7 g |
| Citric acid | 0.6 g |
| $Na_2EDTA$ (pH 8) | 1.12 ml of 0.25 M stock solution |
| $ddH_2O$ | to 1 L. |

###### Additional stock solutions

| **Reagent** | **Formulation** |
|---|---|
| **0.25 M $Na_2EDTA$ (pH 8)** | 9.3 g in 100 mL, adjust to pH 8.0 |

| | |
|---|---|
| **Iron stock** | 1 g Ferric citrate in 100 mL (~40 mM), Requires heat and time to dissolve |
| **Trace metal mix stock** | 0.3 g $H_3BO_3$ (~45 mM), 0.18 g $MnCl_2.4H_20$ (~9 mM), 22 mg $ZnSO_4.7H_20$ (~750 µM), 39 mg $Na_2MoO_4.2H_20$ (~1.6 mM), 8 mg $CuSO_4.5H_20$ (~300 µM), 5mg $Co(NO_3)_2.6H_2O$ (~170 µM) in 100 mL. |
| **$NaHCO_3$ stock** | 8.4 g in 100 mL (~100 mM), filter sterilise |
| **$Na_2CO_3$ stock** | 2 g in 100 mL (~ 190 mM), autoclave |
| **Phosphate stock** | 4 g $K_2HPO_4.3H_20$ in 100 mL (~175 mM), autoclave |
| **Vitamin B12 stock** | 200 mg Cyanocobalamin in 50 mL (4 mg/mL, ~3 mM) |
| **HEPES stock** | 119 g in 500 mL (~1 M), pH to 8.2, autoclave |
| **TES buffer stock** | 23 g in 100 mL (~100 mM), pH to 8.2 |

##### BG-11 liquid medium (1L)

| **Reagent** | **Quantity for 1 L of BG-11** |
|---|---|
| 100x BG-11 | 10 mL |
| Iron stock | 1 mL |
| Trace metal stock | 1 mL |
| Vitamin B12 stock | 100 µL |
| $ddH_2O$ | Up to 978mL |
| $NaHCO_3$ stock | 10 mL: Add after autoclaving base mix |
| HEPES stock | 10 mL: Add after autoclaving base mix |
| $Na_2CO_3$ stock | 1 mL: Add after autoclaving base mix |
| Phosphate stock | 1 mL: Add after autoclaving base mix |

1. Mix 10 mL 100x BG-11, 1 mL Iron Stock, 1 mL Trace metal stock, 100 uL Vitamin B12 stock and add water up to a total of 978 mL (as above)
2. Autoclave
3. Before use, add 10 mL $NaHCO_3$ stock, 10 mL HEPES stock, 1mL $Na_2CO_3$ stock, 1mL phosphate stock

##### BG11 agar plates (1 L)

1. In advance make
   - Agar: 15 g agar in 700 mL water
   - BG-11 plate mixture

| **Reagent** | **Quantity for 1 L of BG-11** |
|---|---|
| 100x BG-11 | 10 mL |
| Iron stock | 1 mL |

| | |
|---|---|
| Trace metal stock | 1 mL |
| Vitamin B12 stock | 100 µL |
| TES buffer stock | 10 mL |
| $Na_2S_2O_3$ | 3 g |
| $ddH_2O$ | Up to 288mL |

2. Autoclave both solutions
3. Before use, melt agar mix and allow to cool
4. Per 100 mL needed, combine use 70 mL molten agar, 29 mL BG-11 plate mixture, 1 mL $NaHCO_3$ stock, 100uL $Na_2CO_3$ stock, 100 uL phosphate stock
5. Add antibiotics at the relevant concentrations and pour plates

#### Affinity purification buffers

##### 2x AP buffer

Adjust to pH 6.8. Store at 4°C

| Reagent | Quantity for 1 L of 2× AP buffer |
|---|---|
| 100 mM HEPES (MW 238.3) | 28.83 g |
| 100 mM KOAc (MW 98.14) | 9.814 g |
| 4 mM $Mg(OAc)_2.4H_2O$ (MW 214.45) | 4 mL of 1 M $Mg(OAc)_2$ (10.725 g in 50 mL $H_2O$) |
| 2 mM $CaCl_2$ | 20 mL of 0.1 M $CaCl_2$ (7.36 g in 500 mL $H_2O$) |
| 400 mM sorbitol (MW 182.17) | 72.87 g |
| $ddH_2O$ | to 900 ml, adjust pH to 6.8 using KOH, top up to 1 L. |

##### 2x AP buffer + PIs

Make fresh on day of AP, sufficient for 12 APs.

| Reagent | Quantity for 25mL |
|---|---|
| 2× AP buffer | 24.5 mL |
| 1 mM NaF (MW 41.99) (Ser/Thr Phosphatase inhibitor) | 50 µL of 1 M solution (0.4199 g in 10 mL H2O) |
| 0.3 mM $Na_3VO_4$ (MW 183.91) (Tyr Phosphatase inhibitor) | 50 µL of 0.3 M solution (0.552 g in 10 mL $H_2O$ adjust pH to 10, boil, readjust pH to 10 and aliquot) |
| Roche cOmplete EDTA-free protease inhibitors | 1 tablet |
| 1 mM PMSF (MW 174.2) (Protease inhibitor) | 500 µL of 100 mM solution (3.484 g in 20 mL isopropanol) |

##### Working AP buffers

- **17mL AP buffer + PIs**: 8.5 mL 2× AP buffer + PIs, 8.5 mL $ddH_2O$

- **30 mL AP buffer + PIs** + 0.1% digitonin: 15 mL 2× AP buffer + PIs, 300 µL 10% digitonin, 14.7 mL dd$H_2O$
- **3mL AP buffer + PIs** + 2% digitonin: 1.5 mL 2× AP buffer + PIs, 600 µL 10% digitonin, 900 µL dd$H_2O$
- Make these fresh on the day of AP, sufficient for 12 APs.

### LB24

- **Per 1 L**: Mix 10 g tryptone, 24 g yeast extract, 5 g NaCl, 1 mL 1 M NaOH, autoclave.

#### A2: Plasmid Sequences and Maps

##### pLM433

>pLM433_CyanoTag_BspQI_BasePlasmid Plasmid used for creating CyanoTag vectors by Golden Gate cloning (BspQI version)
ATGGTTAGTAAGGGTGAGGAAGACAATATGGCAAGCCTGCCGGCAACGCACGAACTGCATATCTTTGGTTCGATCAACGGC
GTGGACTTTGACATGGTCGGCCAGGGGACTGGTAACCCGAACGATGGCTACGAGGAACTGAACCTGAAATCGACAAAAGG
CGATCTGCAATTCTCGCCATGGATTCTCGTGCCGCATATTGGGTACGGCTTTCACCAGTATTTGCCGTATCCAGATGGCATGA
GTCCGTTTCAAGCCGCTATGGTCGATGGTAGCGGCTATCAGGTGCATCGCACTATGCAGTTTGAAGACGGCGCCAGTTTGA
CAGTCAATTACCGTTACACTTATGAAGGCTCGCATATTAAAGGTGAGGCGCAAGTTAAGGGTACCGGGTTTCCCGCCGACGG
TCCCGTGATGACTAATAGCCTGACCGCGGCTGATTGGTGCCGTAGCAAAAAAACCTACCCGAACGACAAAACCATCATTTCC
ACGTTTAAATGGAGCTATACTACAGGTAACGGGAAGCGCTATCGCTCGACGGCGCGCACTACATACACGTTCGCGAAACCG
ATGGCCGCGAATTACCTCAAAAACCAGCCGATGTATGTGTTTCGTAAAACCGAGCTGAAACATAGCAAGACAGAACTGAACT
TTAAAGAGTGGCAGAAAGCATTCACAGACGTCATGGGCATGGATGAACTGTACAAGGGCGGTGGCGATTACAAAGATCACG
ATGGCGATTACAAAGATCACGACATCGACTATAAGGATGACGATGATAAGTAGGCTTCAAATAAAACGAAAGGCTCAGTCGAA
AGACTGGGCCTTTCGTTTTATCTGTTTTTGTCGGTGAACGCTCTCTACTAGAGTCACACTGGCTCACCTTCGGGTGGGCCTT
TCTGCGGGAATTCGATTGATCCGTCGACCTGCAGGGGGGGGGGGGGAAAGCCACGTTGTGTCTCAAAATCTCTGATGTTACA
TTGCACAAGATAAAAATATATCATCATGAACAATAAAACTGTCTGCTTACATAAACAGTAATACAAGGGGTGTTATGAGCCATAT
TCAACGGGAAACGTCTTGCTCGAGGCCGCGATTAAATTCCAACATGGATGCTGATTTATATGGGTATAAATGGGCTCGCGATA
ATGTCGGGCAATCAGGTGCGACAATCTATCGATTGTATGGGAAGCCCGATGCGCCAGAGTTGTTTCTGAAACATGGCAAAG
GTAGCGTTGCCAATGATGTTACAGATGAGATGGTCAGACTAAACTGGCTGACGGAATTTATGCCTCTTCCGACCATCAAGCAT
TTTATCCGTACTCCTGATGATGCATGGTTACTCACCACTGCGATCCCCGGGAAAACAGCATTCCAGGTATTAGAAGAATATCC
TGATTCAGGTGAAAATATTGTTGATGCGCTGGCAGTGTTCCTGCGCCGGTTGCATTCGATTCCTGTTTGTAATTGTCCTTTTA
ACAGCGATCGCGTATTTCGTCTCGCTCAGGCGCAATCACGAATGAATAACGGTTTGGTTGATGCGAGTGATTTTGATGACGA
GCGTAATGGCTGGCCTGTTGAACAAGTCTGGAAAGAAATGCATAAGCTTTTGCCATTCTCACCGGATTCAGTCGTCACTCAT
GGTGATTTCTCACTTGATAACCTTATTTTTGACGAGGGGAAATTAATAGGTTGTATTGATGTTGGACGCGTCGGAATCGCAGA
CCGATACCAGGATCTTGCCATCCTATGGAACTGCCTCGGTGAGTTTTCTCCTTCATTACAGAAACGGCTTTTTCAAAAATATG
GTATTGATAATCCTGATATGAATAAATTGCAGTTTCATTTGATGCTCGATGAGTTTTTCTAATCAGAATTGGTTAATTGGTTGTAA
CACTGGCAGAGCATTACGCTGACTTGACGGGACGGCGGCTTTGTTGAATAAATCGAACTTTTGCTGAGTTGAAGGATCAGAT
CACGCATCTTCCCGACAACGCAGACCGTTCCGTGGCAAAGCAAAAGTTCAAAATCACCAACTGGTCCACCTACAACAAAGC
TCTCATCAACCGTGGCTCCCTCACTTTCTGGCTGGATGATGGGGCGATTCAGGCCTGGTATGAGCCAGCAACACCTTCTTC
ACGAGGCAGACCTCAGCGCTCCTCCACCGCTGCAGTTCACTTACACCGCTTCTCAACCCGGTACGCACCAGAAAATCATTG
ATATGGCCATGAATGGCGTTGGATGCCGGGCAACAGCCCGCATTATGGGCGTTGGCCTCAACACGATTTTACGTCACTTAAA
AAACTCAGGCCGCAGTCGGTAACCTCGCGCATACAGCCGGGCAGTGACGTCATCGTCTGCGCGGAAATGGACGAACAGTG
GGGCTATGTCGGGGCTAAATCGCGCCAGCGCTGGCTGTTTTACGCGTATGACAGTCTCCGGAAGACGGTTGTTGCGCACG
TATTCGGTGAACGCACTATGGCGACGCTGGGGCGTCTTATGAGCCTGCTGTCACCCTTTGACGTGGTGATATGGATGACGG
ATGGCTGGCCGCTGTATGAATCCCGCCTGAAGGGAAAGCTGCACGTAATCAGCAAGCGATATACGCAGCGAATTGAGCGGC
ATAACCTGAATCTGAGGCAGCACCTGGCACGGCTGGGACGGAAGTCGCTGTCGTTCTCAAAATCGGTGGAGCTGCATGAC
AAAGTCATCGGGCATTATCTGAACATAAAACACTATCAATAAGTTGGAATCATTACCAAAAGGTTAGGAATACGGTTAGCCATT
TGCCTGCTTTTATATAGTTCATATGGGATTCACCTTTATGTTGATAAGAAATAAAAGAAAATGCCAATAGGATATCGGCATTTTCT
TTTGCGTTTTCAACGTTTGTAATCGATGGCTTCTGGCTGCTCCAGATATACGGTGGTTTGTGCCGGTTGTGTGCTGGCAATC
ACCTTGCCGCCACGTACCGAATAACGTACCGGAACCTGACGGCGCAGCGCATCAAACCCATTTTCAGCCGGCAGGATAATC
AGGTTGGCGCTGTTTCCGGCGGCAATGCCGTAATCCTGCAAATTCAACGTCCTTGCGCTGTGGTGGGTGATTAAATTCAGG
CCATCGTTAATCTGCCCGTAGCCCATCAACTGGCAAACATGCAGCCCCATATGCAGCACTTGCAGCATATTCGCCGTTCCCA
GCGGATACCACGGATCGAAGACATCATCGTGACCAAAGCAGACGTTAATGCCCGATTCCAGCATCTCTTTAACGCGCGTGAT
GCCGCGACGTTTTGGATACGTATCGAAACGTCCTTGCAGATGAATATTGACCAGCGGGTTGGCGACAAAGTTAATACCGGAC
ATTTTCAGCAAGCGGAACAGGCGTGAGGTATACGCCCCGTTATAGGAGTGCATTGCCGTGGTGTGGCTGGCGGTGACGCG
CGCGCCCATGCCTTCATGGTGCGCCAGGGCAGCAACGGTTTCGACAAAGCGCGACTGCTCGTCATCGATCTCATCACAGT
GAACGTCGATGAGACGGTCGTATTTTTGCGCCAGGGCGAAGGTTTTATGCAGCGATTCCACGCCGTATTCACGGGTAAATTC

AAAATGCGGAATCGCCCCCACTACATCTGCCCCTAAGCGTAACGCCTCTTCCAGCAACGCTTCACCGTTGGGATACGACAA
AATCCCTTCCTGAGGGAAGGCGACGATTTGCAGATCAATCCACGGCGCGACTTCCTGCTTCACTTCCAGCATTGCTTTCAG
CGCAGTTAGCGTTGCATCCGAAACATCGACATGGGTACGCACATGCTGAATGCCGTTGGCAATCTGCCATTTCAGCGTTTGC
CATGCGCGTTGTTTCACATCGTCATGGGTTAATAACGCTTTGCGCTCGGCCCAGCGTTCAATGCCTTCAAACAGCGTGCCG
GACTGATTCCAGTTCGGTTGTCCGGCGGTTTGCGTGGTGTCCAGGTGAATATGTGGCTCCACAAACGGCGGTATAACTAAA
CCTTGTTCGGCATCCAGGCTGTTTTCAGTTATGGGCATCACGCCGGATTGCGCATCAATGGCGCTGATTTTTCCGTCCTGCA
GATGAATCTGCCACAGCCCCTCTTCGCCTGGTAACCGGGCGTTAATAATTGTTTGTAAAGCGTTATTCGACATCGTTCATGTC
TCCTTTTTTATGTACTGTGTTAGCGGTCTGCTTCTTCCAGCCCTCCTGTTTGAAGATGGCAAGTTAGTTACGCACAATAAAAAA
AGACCTAAAATATGTAAGGGGTGACGCCAAAGTATACACTTTGCCCTTTACACATTTTAGGTCTTGCCTGCTTTATCAGTAACA
AACCCGCGCGATTTACTTTTCGACCTCATTCTATTAGATTCTCGTTTGGATTGCAACTGGTCTATTTTCCTCTTTTGTTTGATAG
AAAATCATAAAAGGATTTGCAGACTACGGGCCTAAAGGTTAGTAAGGGTGAGGAAGACAATATGGCAAGCCTGCCGGCAAC
GCACGAACTGCATATCTTTGGTTCGATCAACGGCGTGGACTTTGACATGGTCGGCCAGGGGACTGGTAACCCGAACGATGG
CTACGAGGAACTGAACCTGAAATCGACAAAAGGCGATCTGCAATTCTCGCCATGGATTCTCGTGCCGCATATTGGGTACGGC
TTTCACCAGTATTTGCCGTATCCAGATGGCATGAGTCCGTTTCAAGCCGCTATGGTCGATGGTAGCGGCTATCAGGTGCATC
GCACTATGCAGTTTGAAGACGGCGCCAGTTTGACAGTCAATTACCGTTACACTTATGAAGGCTCGCATATTAAAGGTGAGGC
GCAAGTTAAGGGTACCGGGTTTCCCGCCGACGGTCCCGTGATGACTAATAGCCTGACCGCGGCTGATTGGTGCCGTAGCA
AAAAAACCTACCCGAACGACAAAACCATCATTTCCACGTTTAAATGGAGCTATACTACAGGTAACGGGAAGCGCTATCGCTCG
ACGGCGCGCACTACATACACGTTCGCGAAACCGATGGCCGCGAATTACCTCAAAAACCAGCCGATGTATGTGTTTCGTAAAA
CCGAGCTGAAACATAGCAAGACAGAACTGAACTTTAAAGAGTGGCAGAAAGCATTCACAGACGTCATGGGCATGGATGAAC
TGTACAAGGGCGGTGGCGATTACAAAGATCACGATGGCGATTACAAAGATCACGACATCGACTATAAGGATGACGATGATAA
GTAGTGAAGAGCTAAAAGCCAGATAACAGTATGCGTATTTGCGCGCTGATTTTTGCGGTATAAGAATATATACTGATATGTATAC
CCGAAGTATGTCAAAAAGAGGTATGCTATGAAGCAGCGTATTACAGTGACAGTTGACAGCGACAGCTATCAGTTGCTCAAGG
CATATATGATGTCAATATCTCCGGTCTGGTAAGCACAACCATGCAGAATGAAGCCCGTCGTCTGCGTGCCGAACGCTGGAAA
GCGGAAAATCAGGAAGGGATGGCTGAGGTCGCCCGGTTTATTGAAATGAACGGCTCTTTTGCTGACGAGAACAGGGGCTG
GTGAAATGCAGTTTAAGGTTTACACCTATAAAAGAGAGAGCCGTTATCGTCTGTTTGTGGATGTACAGAGTGATATTATTGACA
CGCCCGGGCGACGGATGGTGATCCCCCTGGCCAGTGCACGTCTGCTGTCAGATAAAGTCCCCCGTGAACTTTACCCGGTG
GTGCATATCGGGGATGAAAGCTGGCGCATGATGACCACCGATATGGCCAGTGTGCCGGTGTCCGTTATCGGGGAAGAAGTG
GCTGATCTCAGCCACCGCGAAAATGACATCAAAAACGCCATTAACCTGATGTTCTGGGGAATATAAATGTCAGGCTCCCTTAT
ACACAGCCAGTCTGCAGGTCGACCATAGTGCTCTTCACTTGAGACTCTTTCCATAGGCTCCGCCCCCCTGACGAGCATCAC
AAAAATCGACGCTCAAGTCAGAGGTGGCGAAACCCGACAGGACTATAAAGATACCAGGCGTTTCCCCCTGGAAGCTCCCTC
GTGCGCTCTCCTGTTCCGACCCTGCCGCTTACCGGATACCTGTCCGCCTTTCTCCCTTCGGGAAGCGTGGCGCTTTCTCAT
AGCTCACGCTGTAGGTATCTCAGTTCGGTGTAGGTCGTTCGCTCCAAGCTGGGCTGTGTGCACGAACCCCCCGTTCAGCC
CGACCGCTGCGCCTTATCCGGTAACTATCGTCTTGAGCCCAACCCGGTAAGACACGACTTATCGCCACTGGCAGCAGCCAC
TGGTAACAGGATTAGCAGAGCGAGGTATGTAGGCGGTGCTACAGAGTTCTTGAAGTGGTGGCCTAACTACGGCTACACTAG
AAGAACAGTATTTGGTATCTGCGCTCTGCTGAAGCCAGTTACCTTCGGAAAAAGAGTTGGTAGCTCTTGATCCGGCAAACAA
ACCACCGCTGGTAGCGGTGGTTTTTTTGTTTGCAAGCAGCAGATTACGCGCAGAAAAAAAGGATCTCAAGAAGATCCTTTGA
TCTTTTCTACGGGGTCTGACGCTCAGTGGAACGAAAACTCACGTTAAGGGATTTTGGTCATGAGATTATCAAAAAGGATCTTC
ACCTAGATCCTTTTAAATTAAAAATGAAGTTTTAAATCAATCTAAAGTATATATGAGTAAACTTGGTCTGACAGTTACCAATGCTT
AATCAGTGAGGCACCTATCTCAGCGATCTGTCTATTTCGTTCATCCATAGTTGCCTGGCTCCCCGTCGTGTAGATAACTACGA
TACGGGAGGGCTTACCATCTGGCCCCAGTGCTGCAATGATACCGCGTGACCCACGCTCACCGGCTCCAGATTTATCAGCAA
TAAACCAGCCAGCCGGAAGGGCCGAGCGCAGAAGTGGTCCTGCAACTTTATCCGCCTCCATCCAGTCTATTAATTGTTGCC
GGGAAGCTAGAGTAAGTAGTTCGCCAGTTAATAGTTTGCGCAACGTTGTTGCCATTGCTACAGGCATCGTGGTGTCACGCTC
GTCGTTTGGTATGGCTTCATTCAGCTCCGGTTCCCAACGATCAAGGCGAGTTACATGATCCCCCATGTTGTGCAAAAAAGCG
GTTAGCTCCTTCGGTCCTCCGATCGTTGTCAGAAGTAAGTTGGCCGCAGTGTTATCACTCATGGTTATGGCAGCACTGCATA
ATTCTCTTACTGTCATGCCATCCGTAAGATGCTTTTCTGTGACTGGTGAGTACTCAACCAAGTCATTCTGAGAATAGTGTATGC
GGCGACCGAGTTGCTCTTGCCCGGCGTCAATACGGGATAATACCGCGCCACATAGCAGAACTTTAAAAGTGCTCATCATTGG
AAAACGTTCTTCGGGGCGAAAACTCTCAAGGATCTTACCGCTGTTGAGATCCAGTTCGATGTAACCCACTCGTGCACCCAAC
TGATCTTCAGCATCTTTTACTTTCACCAGCGTTTCTGGGTGAGCAAAAACAGGAAGGCAAAATGCCGCAAAAAAGGGAATAA
GGGCGACACGGAAATGTTGAATACTCATACTCTTCCTTTTTCAATATTATTGAAGCATTTATCAGGGTTATTGTCTCATGAGCG
GATACATATTTGAATGTATTTAGAAAAATAAACAAATAGGGGTTCCGCGGAGTCAGACTTGAAGAGCTAAAAGCCAGATAACA
GTATGCGTATTTGCGCGCTGATTTTTGCGGTATAAGAATATATACTGATATGTATACCCGAAGTATGTCAAAAAGAGGTATGCTA
TGAAGCAGCGTATTACAGTGACAGTTGACAGCGACAGCTATCAGTTGCTCAAGGCATATATGATGTCAATATCTCCGGTCTGG
TAAGCACAACCATGCAGAATGAAGCCCGTCGTCTGCGTGCCGAACGCTGGAAAGCGGAAAATCAGGAAGGGATGGCTGAG
GTCGCCCGGTTTATTGAAATGAACGGCTCTTTTGCTGACGAGAACAGGGGCTGGTGAAATGCAGTTTAAGGTTTACACCTAT
AAAAGAGAGAGCCGTTATCGTCTGTTTGTGGATGTACAGAGTGATATTATTGACACGCCCGGGCGACGGATGGTGATCCCCC
TGGCCAGTGCACGTCTGCTGTCAGATAAAGTCCCCCGTGAACTTTACCCGGTGGTGCATATCGGGGATGAAAGCTGGCGCA
TGATGACCACCGATATGGCCAGTGTGCCGGTTTCCGTTATCGGGGAAGAAGTGGCTGATCTCAGCCACCGCGAAAATGACA
TCAAAAACGCCATTAACCTGATGTTCTGGGGAATATAAATGTCAGGCTCCCTTATACACAGCCAGTCTGCAGGTCGACCATAG
TGCTCTTCAGGCGATCTGGGTGGCAGCGGCGGCCGC

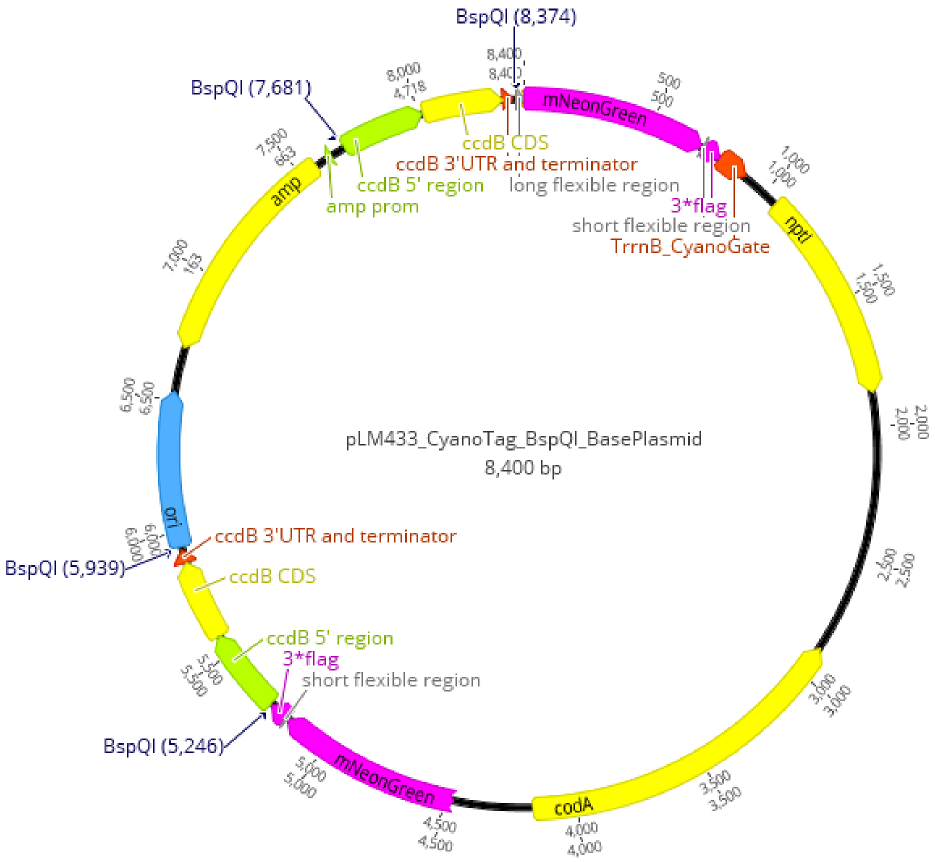


### pLM434

>pLM434_CyanoTag_BsaI_BasePlasmid Plasmid used for creating CyanoTag vectors by Golden Gate cloning (BsaI version)
ATGGTTAGTAAGGGTGAGGAAGACAATATGGCAAGCCTGCCGGCAACGCACGAACTGCATATCTTTGGTTCGATCAACGGC
GTGGACTTTGACATGGTCGGCCAGGGGACTGGTAACCCGAACGATGGCTACGAGGAACTGAACCTGAAATCGACAAAAGG
CGATCTGCAATTCTCGCCATGGATTCTCGTGCCGCATATTGGGTACGGCTTTCACCAGTATTTGCCGTATCCAGATGGCATGA
GTCCGTTTCAAGCCGCTATGGTCGATGGTAGCGGCTATCAGGTGCATCGCACTATGCAGTTTGAAGACGGCGCCAGTTTGA
CAGTCAATTACCGTTACACTTATGAAGGCTCGCATATTAAAGGTGAGGCGCAAGTTAAGGGTACCGGGTTTCCCGCCGACGG
TCCCGTGATGACTAATAGCCTGACCGCGGCTGATTGGTGCCGTAGCAAAAAAACCTACCCGAACGACAAAACCATCATTTCC
ACGTTTAAATGGAGCTATACTACAGGTAACGGGAAGCGCTATCGCTCGACGGCGCGCACTACATACACGTTCGCGAAACCG
ATGGCCGCGAATTACCTCAAAAACCAGCCGATGTATGTGTTTCGTAAAACCGAGCTGAAACATAGCAAGACAGAACTGAACT
TTAAAGAGTGGCAGAAAGCATTCACAGACGTCATGGGCATGGATGAACTGTACAAGGGCGGTGGCGATTACAAAGATCACG
ATGGCGATTACAAAGATCACGACATCGACTATAAGGATGACGATGATAAGTAGGCTTCAAATAAAACGAAAGGCTCAGTCGAA
AGACTGGGCCTTTCGTTTTATCTGTTTTTGTCGGTGAACGCTCTCTACTAGAGTCACACTGGCTCACCTTCGGGTGGGCCTT
TCTGCGGGAATTCGATTGATCCGTCGACCTGCAGGGGGGGGGGGGGAAAGCCACGTTGTGTCTCAAAATCTCTGATGTTACA
TTGCACAAGATAAAAATATATCATCATGAACAATAAAACTGTCTGCTTACATAAACAGTAATACAAGGGGTGTTATGAGCCATAT
TCAACGGGAAACGTCTTGCTCGAGGCCGCGATTAAATTCCAACATGGATGCTGATTTATATGGGTATAAATGGGCTCGCGATA
ATGTCGGGCAATCAGGTGCGACAATCTATCGATTGTATGGGAAGCCCGATGCGCCAGAGTTGTTTCTGAAACATGGCAAAG
GTAGCGTTGCCAATGATGTTACAGATGAGATGGTCAGACTAAACTGGCTGACGGAATTTATGCCTCTTCCGACCATCAAGCAT
TTTATCCGTACTCCTGATGATGCATGGTTACTCACCACTGCGATCCCCGGGAAAACAGCATTCCAGGTATTAGAAGAATATCC
TGATTCAGGTGAAAATATTGTTGATGCGCTGGCAGTGTTCCTGCGCCGGTTGCATTCGATTCCTGTTTGTAATTGTCCTTTTA
ACAGCGATCGCGTATTTCGTCTCGCTCAGGCGCAATCACGAATGAATAACGGTTTGGTTGATGCGAGTGATTTTGATGACGA
GCGTAATGGCTGGCCTGTTGAACAAGTCTGGAAAGAAATGCATAAGCTTTTGCCATTCTCACCGGATTCAGTCGTCACTCAT
GGTGATTTCTCACTTGATAACCTTATTTTTGACGAGGGGAAATTAATAGGTTGTATTGATGTTGGACGCGTCGGAATCGCAGA
CCGATACCAGGATCTTGCCATCCTATGGAACTGCCTCGGTGAGTTTTCTCCTTCATTACAGAAACGGCTTTTTCAAAAATATG
GTATTGATAATCCTGATATGAATAAATTGCAGTTTCATTTGATGCTCGATGAGTTTTTCTAATCAGAATTGGTTAATTGGTTGTAA
CACTGGCAGAGCATTACGCTGACTTGACGGGACGGCGGCTTTGTTGAATAAATCGAACTTTTGCTGAGTTGAAGGATCAGAT
CACGCATCTTCCCGACAACGCAGACCGTTCCGTGGCAAAGCAAAAGTTCAAAATCACCAACTGGTCCACCTACAACAAAGC
TCTCATCAACCGTGGCTCCCTCACTTTCTGGCTGGATGATGGGGCGATTCAGGCCTGGTATGAGCCAGCAACACCTTCTTC
ACGAGGCAGACCTCAGCGCTCCTCCACCGCTGCAGTTCACTTACACCGCTTCTCAACCCGGTACGCACCAGAAAATCATTG
ATATGGCCATGAATGGCGTTGGATGCCGGGCAACAGCCCGCATTATGGGCGTTGGCCTCAACACGATTTTACGTCACTTAAA
AAACTCAGGCCGCAGTCGGTAACCTCGCGCATACAGCCGGGCAGTGACGTCATCGTCTGCGCGGAAATGGACGAACAGTG

GGGCTATGTCGGGGCTAAATCGCGCCAGCGCTGGCTGTTTTACGCGTATGACAGTCTCCGGAAGACGGTTGTTGCGCACG
TATTCGGTGAACGCACTATGGCGACGCTGGGGCGTCTTATGAGCCTGCTGTCACCCTTTGACGTGGTGATATGGATGACGG
ATGGCTGGCCGCTGTATGAATCCCGCCTGAAGGGAAAGCTGCACGTAATCAGCAAGCGATATACGCAGCGAATTGAGCGGC
ATAACCTGAATCTGAGGCAGCACCTGGCACGGCTGGGACGGAAGTCGCTGTCGTTCTCAAAATCGGTGGAGCTGCATGAC
AAAGTCATCGGGCATTATCTGAACATAAAACACTATCAATAAGTTGGAATCATTACCAAAAGGTTAGGAATACGGTTAGCCATT
TGCCTGCTTTTATATAGTTCATATGGGATTCACCTTTATGTTGATAAGAAATAAAAGAAAATGCCAATAGGATATCGGCATTTTCT
TTTGCGTTTTCAACGTTTGTAATCGATGGCTTCTGGCTGCTCCAGATATACGGTGGTTTGTGCCGGTTGTGTGCTGGCAATC
ACCTTGCCGCCACGTACCGAATAACGTACCGGAACCTGACGGCGCAGCGCATCAAACCCATTTTCAGCCGGCAGGATAATC
AGGTTGGCGCTGTTTCCGGCGGCAATGCCGTAATCCTGCAAATTCAACGTCCTTGCGCTGTGGTGGGTGATTAAATTCAGG
CCATCGTTAATCTGCCCGTAGCCCATCAACTGGCAAACATGCAGCCCCATATGCAGCACTTGCAGCATATTCGCCGTTCCCA
GCGGATACCACGGATCGAAGACATCATCGTGACCAAAGCAGACGTTAATGCCCGATTCCAGCATCTCTTTAACGCGCGTGAT
GCCGCGACGTTTTGGATACGTATCGAAACGTCCTTGCAGATGAATATTGACCAGCGGGTTGGCGACAAAGTTAATACCGGAC
ATTTTCAGCAAGCGGAACAGGCGTGAGGTATACGCCCCGTTATAGGAGTGCATTGCCGTGGTGTGGCTGGCGGTGACGCG
CGCGCCCATGCCTTCATGGTGCGCCAGGGCAGCAACGGTTTCGACAAAGCGCGACTGCTCGTCATCGATCTCATCACAGT
GAACGTCGATGAGACGGTCGTATTTTTGCGCCAGGGCGAAGGTTTTATGCAGCGATTCCACGCCGTATTCACGGGTAAATTC
AAAATGCGGAATCGCCCCCACTACATCTGCCCCTAAGCGTAACGCCTCTTCCAGCAACGCTTCACCGTTGGGATACGACAA
AATCCCTTCCTGAGGGAAGGCGACGATTTGCAGATCAATCCACGGCGCGACTTCCTGCTTCACTTCCAGCATTGCTTTCAG
CGCAGTTAGCGTTGCATCCGAAACATCGACATGGGTACGCACATGCTGAATGCCGTTGGCAATCTGCCATTTCAGCGTTTGC
CATGCGCGTTGTTTCACATCGTCATGGGTTAATAACGCTTTGCGCTCGGCCCAGCGTTCAATGCCTTCAAACAGCGTGCCG
GACTGATTCCAGTTCGGTTGTCCGGCGGTTTGCGTGGTGTCCAGGTGAATATGTGGCTCCACAAACGGCGGTATAACTAAA
CCTTGTTCGGCATCCAGGCTGTTTTCAGTTATGGGCATCACGCCGGATTGCGCATCAATGGCGCTGATTTTTCCGTCCTGCA
GATGAATCTGCCACAGCCCCTCTTCGCCTGGTAACCGGGCGTTAATAATTGTTTGTAAAGCGTTATTCGACATCGTTCATGTC
TCCTTTTTTATGTACTGTGTTAGCGGTCTGCTTCTTCCAGCCCTCCTGTTTGAAGATGGCAAGTTAGTTACGCACAATAAAAAA
AGACCTAAAATATGTAAGGGGTGACGCCAAAGTATACACTTTGCCCTTTACACATTTTAGGTCTTGCCTGCTTTATCAGTAACA
AACCCGCGCGATTTACTTTTCGACCTCATTCTATTAGATTCTCGTTTGGATTGCAACTGGTCTATTTTCCTCTTTTGTTTGATAG
AAAATCATAAAAGGATTTGCAGACTACGGGCCTAAAGGTTAGTAAGGGTGAGGAAGACAATATGGCAAGCCTGCCGGCAAC
GCACGAACTGCATATCTTTGGTTCGATCAACGGCGTGGACTTTGACATGGTCGGCCAGGGGACTGGTAACCCGAACGATGG
CTACGAGGAACTGAACCTGAAATCGACAAAAGGCGATCTGCAATTCTCGCCATGGATTCTCGTGCCGCATATTGGGTACGGC
TTTCACCAGTATTTGCCGTATCCAGATGGCATGAGTCCGTTTCAAGCCGCTATGGTCGATGGTAGCGGCTATCAGGTGCATC
GCACTATGCAGTTTGAAGACGGCGCCAGTTTGACAGTCAATTACCGTTACACTTATGAAGGCTCGCATATTAAAGGTGAGGC
GCAAGTTAAGGGTACCGGGTTTCCCGCCGACGGTCCCGTGATGACTAATAGCCTGACCGCGGCTGATTGGTGCCGTAGCA
AAAAAACCTACCCGAACGACAAAACCATCATTTCCACGTTTAAATGGAGCTATACTACAGGTAACGGGAAGCGCTATCGCTCG
ACGGCGCGCACTACATACACGTTCGCGAAACCGATGGCCGCGAATTACCTCAAAAACCAGCCGATGTATGTGTTTCGTAAAA
CCGAGCTGAAACATAGCAAGACAGAACTGAACTTTAAAGAGTGGCAGAAAGCATTCACAGACGTCATGGGCATGGATGAAC
TGTACAAGGGCGGTGGCGATTACAAAGATCACGATGGCGATTACAAAGATCACGACATCGACTATAAGGATGACGATGATAA
GTAGTGAGACCTAAAAGCCAGATAACAGTATGCGTATTTGCGCGCTGATTTTTGCGGTATAAGAATATATACTGATATGTATACC
CGAAGTATGTCAAAAAGAGGTATGCTATGAAGCAGCGTATTACAGTGACAGTTGACAGCGACAGCTATCAGTTGCTCAAGGC
ATATATGATGTCAATATCTCCGGTCTGGTAAGCACAACCATGCAGAATGAAGCCCGTCGTCTGCGTGCCGAACGCTGGAAAG
CGGAAAATCAGGAAGGGATGGCTGAGGTCGCCCGGTTTATTGAAATGAACGGCTCTTTTGCTGACGAGAACAGGGGCTGG
TGAAATGCAGTTTAAGGTTTACACCTATAAAAGAGAGAGCCGTTATCGTCTGTTTGTGGATGTACAGAGTGATATTATTGACAC
GCCCGGGCGACGGATGGTGATCCCCCTGGCCAGTGCACGTCTGCTGTCAGATAAAGTCCCCCGTGAACTTTACCCGGTGG
TGCATATCGGGGATGAAAGCTGGCGCATGATGACCACCGATATGGCCAGTGTGCCGGTTTCCGTTATCGGGGAAGAAGTGG
CTGATCTCAGCCACCGCGAAAATGACATCAAAAACGCCATTAACCTGATGTTCTGGGGAATATAAATGTCAGGCTCCCTTATA
CACAGCCAGTCTGCAGGTCGACCATAGTGGTCTCACTTGAGACTCTTTCCATAGGCTCCGCCCCCCTGACGAGCATCACAA
AAATCGACGCTCAAGTCAGAGGTGGCGAAACCCGACAGGACTATAAAGATACCAGGCGTTTCCCCCTGGAAGCTCCCTCGT
GCGCTCTCCTGTTCCGACCCTGCCGCTTACCGGATACCTGTCCGCCTTTCTCCCTTCGGGAAGCGTGGCGCTTTCTCATAG
CTCACGCTGTAGGTATCTCAGTTCGGTGTAGGTCGTTCGCTCCAAGCTGGGCTGTGTGCACGAACCCCCCGTTCAGCCCG
ACCGCTGCGCCTTATCCGGTAACTATCGTCTTGAGCCCAACCCGGTAAGACACGACTTATCGCCACTGGCAGCAGCCACTG
GTAACAGGATTAGCAGAGCGAGGTATGTAGGCGGTGCTACAGAGTTCTTGAAGTGGTGGCCTAACTACGGCTACACTAGAA
GAACAGTATTTGGTATCTGCGCTCTGCTGAAGCCAGTTACCTTCGGAAAAAGAGTTGGTAGCTCTTGATCCGGCAAACAAAC
CACCGCTGGTAGCGGTGGTTTTTTTGTTTGCAAGCAGCAGATTACGCGCAGAAAAAAAGGATCTCAAGAAGATCCTTTGATC
TTTTCTACGGGGTCTGACGCTCAGTGGAACGAAAACTCACGTTAAGGGATTTTGGTCATGAGATTATCAAAAAGGATCTTCAC
CTAGATCCTTTTAAATTAAAAATGAAGTTTTAAATCAATCTAAAGTATATATGAGTAAACTTGGTCTGACAGTTACCAATGCTTAA
TCAGTGAGGCACCTATCTCAGCGATCTGTCTATTTCGTTCATCCATAGTTGCCTGGCTCCCCGTCGTGTAGATAACTACGATA
CGGGAGGGCTTACCATCTGGCCCCAGTGCTGCAATGATACCGCGTGACCCACGCTCACCGGCTCCAGATTTATCAGCAATA
AACCAGCCAGCCGGAAGGGCCGAGCGCAGAAGTGGTCCTGCAACTTTATCCGCCTCCATCCAGTCTATTAATTGTTGCCGG
GAAGCTAGAGTAAGTAGTTCGCCAGTTAATAGTTTGCGCAACGTTGTTGCCATTGCTACAGGCATCGTGGTGTCACGCTCGT
CGTTTGGTATGGCTTCATTCAGCTCCGGTTCCCAACGATCAAGGCGAGTTACATGATCCCCCATGTTGTGCAAAAAAGCGGT
TAGCTCCTTCGGTCCTCCGATCGTTGTCAGAAGTAAGTTGGCCGCAGTGTTATCACTCATGGTTATGGCAGCACTGCATAATT
CTCTTACTGTCATGCCATCCGTAAGATGCTTTTCTGTGACTGGTGAGTACTCAACCAAGTCATTCTGAGAATAGTGTATGCGG
CGACCGAGTTGCTCTTGCCCGGCGTCAATACGGGATAATACCGCGCCACATAGCAGAACTTTAAAAGTGCTCATCATTGGAA
AACGTTCTTCGGGGCGAAAACTCTCAAGGATCTTACCGCTGTTGAGATCCAGTTCGATGTAACCCACTCGTGCACCCAACT
GATCTTCAGCATCTTTTACTTTCACCAGCGTTTCTGGGTGAGCAAAAACAGGAAGGCAAAATGCCGCAAAAAAGGGAATAAG
GGCGACACGGAAATGTTGAATACTCATACTCTTCCTTTTTCAATATTATTGAAGCATTTATCAGGGTTATTGTCTCATGAGCGG
ATACATATTTGAATGTATTTAGAAAAATAAACAAATAGGGGTTCCGCGGAGTCAGACTTGAGACCTAAAAGCCAGATAACAGTA
TGCGTATTTGCGCGCTGATTTTTGCGGTATAAGAATATATACTGATATGTATACCCGAAGTATGTCAAAAAGAGGTATGCTATGA

AGCAGCGTATTACAGTGACAGTTGACAGCGACAGCTATCAGTTGCTCAAGGCATATATGATGTCAATATCTCCGGTCTGGTAA
GCACAACCATGCAGAATGAAGCCCGTCGTCTGCGTGCCGAACGCTGGAAAGCGGAAAATCAGGAAGGGATGGCTGAGGT
CGCCCGGTTTATTGAAATGAACGGCTCTTTTGCTGACGAGAACAGGGGCTGGTGAAATGCAGTTTAAGGTTTACACCTATAA
AAGAGAGAGCCGTTATCGTCTGTTTGTGGATGTACAGAGTGATATTATTGACACGCCCGGGCGACGGATGGTGATCCCCCT
GGCCAGTGCACGTCTGCTGTCAGATAAAGTCCCCCGTGAACTTTACCCGGTGGTGCATATCGGGGATGAAAGCTGGCGCAT
GATGACCACCGATATGGCCAGTGTGCCGGTTTCCGTTATCGGGGAAGAAGTGGCTGATCTCAGCCACCGCGAAAATGACAT
CAAAAACGCCATTAACCTGATGTTCTGGGGAATATAAATGTCAGGCTCCCTTATACACAGCCAGTCTGCAGGTCGACCATAGT
GGTCTCAGGCGATCTGGGTGGCAGCGGCGGCCGC

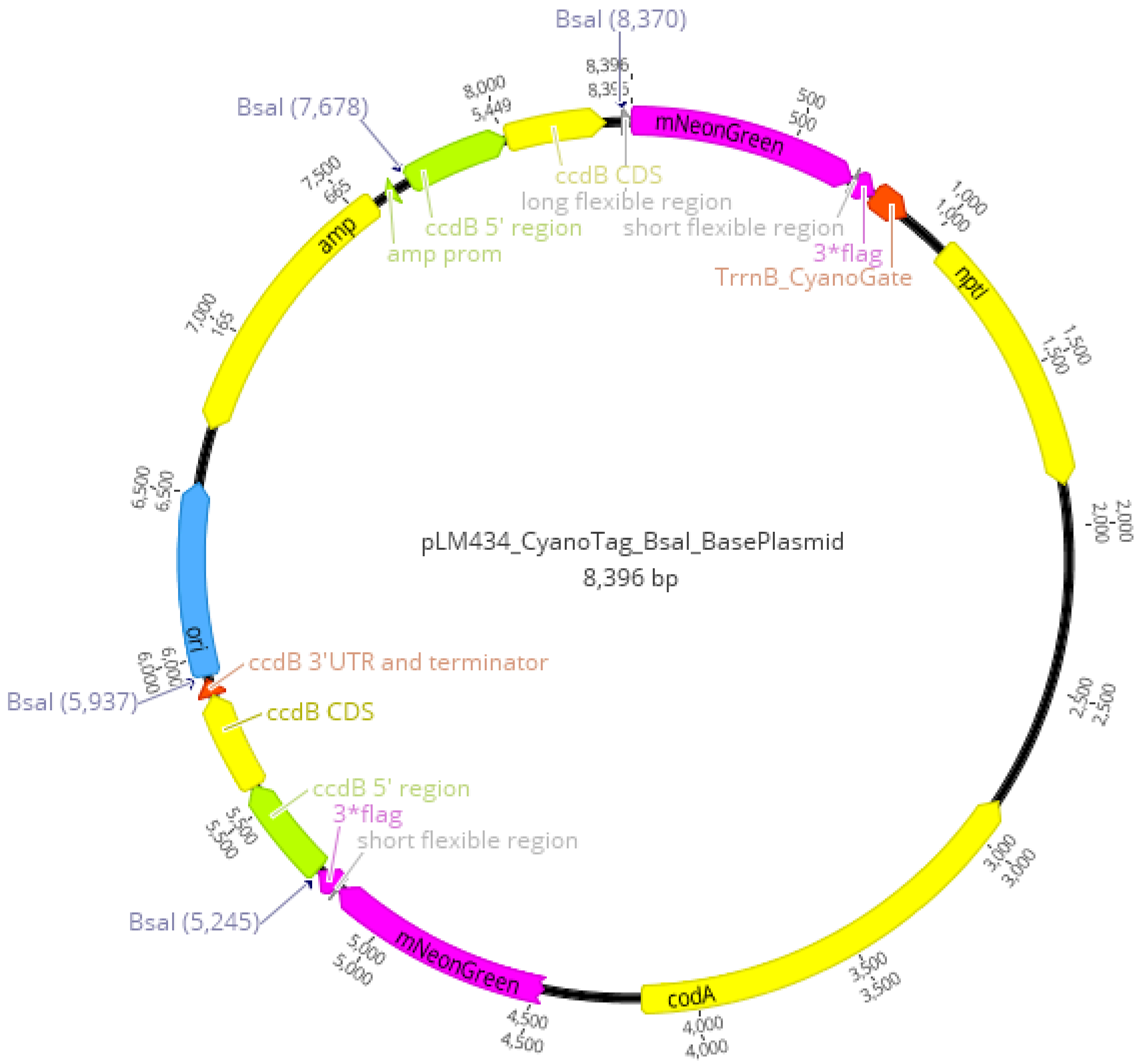


### A3: Macro for making montages from single frames of images from the Elyra 7 in Fiji

*Adapted from a previous Macro supplied by James Barrett*

```
input = getDirectory("Input Directory");
output = getDirectory("Output Directory");
someString = getString("Filetype", ".czi");
suffix = someString
```

```
processFolder(input);

function processFolder(input) {
	list = getFileList(input);
	for (i = 0; i < list.length; i++) {
		if(File.isDirectory(input + list[i]))
			processFolder("" + input + list[i]);
		if(endsWith(list[i], suffix))
			processFile(input, output, list[i]);
		print( list[i] );
	}
}

function processFile(input, output, file) {
	open(input + file);
	name=getTitle;
	selectWindow(name);
	run("Split Channels");
	list2 = getList("image.titles");
	for( i = 0; i < list2.length; i++ ) print( list2[i] );
	run("Merge Channels...", "c6="+list2[0]+" c2="+list2[1]+" create  ignore");
	run("Split Channels");
	run("Merge Channels...", "c2=C1-Composite c6=C2-Composite create keep ignore");
	run("Scale Bar...", "width=5 height=4 font=24 color=White background=None location=[Lower Right] bold overlay");
	run("Flatten");
	selectWindow("C1-Composite");
	run("RGB Color");
	selectWindow("C2-Composite");
	run("RGB Color");
	run("RGB Color", "C2-Composite");
	run("Combine...", "stack1=[C2-Composite] stack2=[C1-Composite]");
	run("Combine...", "stack1=[Combined Stacks] stack2=[Composite (RGB)]");
	saveAs("Tiff", output + file + "_out");
	close("*");
}
```
